# Enriching Representation Learning Using 53 Million Patient Notes through Human Phenotype Ontology Embedding

**DOI:** 10.1101/2022.07.20.500809

**Authors:** Maryam Daniali, Peter D. Galer, David Lewis-Smith, Shridhar Parthasarathy, Edward Kim, Dario D. Salvucci, Jeffrey M. Miller, Scott Haag, Ingo Helbig

## Abstract

The Human Phenotype Ontology (HPO) is a dictionary of more than 15,000 clinical phenotypic terms with defined semantic relationships, developed to standardize their representation for phenotypic analysis. Over the last decade, the HPO has been used to accelerate the implementation of precision medicine into clinical practice. In addition, recent research in representation learning, specifically in graph embedding, has led to notable progress in automated prediction via learned features. Here, we present a novel approach to phenotype representation by incorporating phenotypic frequencies based on 53 million full-text health care notes from more than 1.5 million individuals. We demonstrate the efficacy of our proposed phenotype embedding technique by comparing our work to existing phenotypic similarity-measuring methods. Using phenotype frequencies in our embedding technique, we are able to identify phenotypic similarities that surpass the current computational models. In addition, we show that our embedding technique aligns with domain experts’ judgment at a level that exceeds their agreement. We show that our proposed technique efficiently represents complex and multidimensional phenotypes in HPO format, which can then be used as input for various downstream tasks that require deep phenotyping, including patient similarity analyses and disease trajectory prediction.

## 1 Introduction

Electronic medical records (EMRs) have been implemented in the majority of US hospitals and accumulate clinical data at a massive scale [1]. While initially created for billing purposes [2], EMRs increasingly represent a major source of data in clinical research efforts to improve patient care. However, automated interpretation of EMRs is challenging, particularly for diagnoses that incorporate dynamic and diverse sets of clinical features. Phenotypes, a set of observable characteristics and clinical traits, have an essential role in connecting clinical research and practice. Algorithms that use phenotypes to find similarities and differences between patients play a foundational role in EMR research [3]. However, phenotypic descriptions available in EMRs are often stored in the form of unstructured data and do not allow for direct comparisons.

The Human Phenotype Ontology (HPO) is one approach to overcome these limitations (human-phenotype-ontology.org). The HPO is a standardized representation of more than 15,000 clinical phenotypic concepts and their relationships based on expert knowledge. We have contributed to HPO terminology since 2010 [4-7]. The HPO has been widely used for harmonization of clinical features in various studies, including, but not limited to, semantic unification of common and rare diseases [8], genetic discoveries in pediatric epilepsy [9, 10], and delineation of longitudinal phenotypes [11, 12]. In addition, the HPO is commonly used for genomic studies and allows for analyses of clinical data at a scale that is required by current and future initiatives. For example, large national and international initiatives have started to systematically link biorepositories to EMR data, including up to 80,000 cases and 500,000 controls [13-15], highlighting the possibilities for novel biological insight at scale.

The HPO can be modeled as a directed acyclic graph (DAG) in a computational system, where each phenotype is presented as a node with a unique identifier and is connected to its parent phenotypes by “is a” relationships in the form of directed edges. This structure guarantees that if a disease or gene is annotated to a phenotypic term, it will also be annotated to all its ancestral terms (higher-level concepts within the larger phenotypic tree). The HPO is regularly updated to incorporate advances in phenotypic conceptualization [16].

Extraction of phenotypic concepts is a crucial step in any automated pipeline to exploit the scale of EMR data in clinical research. Natural Language Processing (NLP) pipelines such as cTAKES [17], ClinPhen [18], and MetaMap [19] are commonly used to derive phenotypic concepts from the EMR, effectively allowing the transition from unstructured free text to structured representations. While these NLP pipelines share a common goal, that is, extracting phenotypes from clinical free text, they have different components and employ a variety of procedures, thus, should be evaluated from different endpoints such as precision, sensitivity, ease of use, and speed [18, 20]. Furthermore, phenotype extraction typically serves as only a starting point to more complex analyses. Therefore, studies need to perform additional analyses on the extracted phenotypes and their relationships to accomplish tasks like patient comparisons and predicting patient status [21]. Manual analysis of phenotypes is non-scalable, resource-intensive, and virtually impossible for larger cohorts. Accordingly, reliable algorithms for computational phenotype analysis are urgently needed.

Comparing phenotypes to one another is a common building block for downstream clinical tasks. Methods for measuring phenotypic similarities, using measures such as the Resnik score [22], information coefficient [23], and graph information content [24], show promise for diagnoses and open doors to novel biological insights such as genetic discoveries [9, 25, 26]. However, these methods are not generally transferable to other tasks and require a significant amount of computation, even with minor changes to the data. As a result, while these methods perform relatively well on limited data with hundreds of patients, they are computationally intensive for large-scale data with millions or even thousands of patients with diverse clinical representations [18, 26].

Representation learning is a group of machine learning algorithms that discovers and learns representations of data, making it easier to extract information that can be used for various tasks such as classification and prediction [27]. Recent work in representation learning has shown success in discovering useful representations without relying on procedural techniques, especially in domains with complex and large data [27]. Embedding algorithms, discussed in **Section 2**, are a branch of representation learning that model discrete objects as continuous vectors. They offer a compact representation that captures similarities between the original objects and have revolutionized data processing and analysis in many domains, including text processing, by representing words in a compact space [28, 29]. Embedding algorithms have also been extended to encode other data structures such as nodes and graphs [30, 31]. A few studies have applied representation learning and embedding techniques in the health domain and presented promising results on specific phenotypes for a limited number of diseases [32, 33].

Here, we map 53 million full-text health care notes of more than 1.5 million individuals to HPO terms and perform graph embedding to assess the possibilities and limitations of representation learning techniques. We demonstrate that phenotype embedding, in some scenarios, can exceed more computationally intense measures of phenotype similarity, providing a framework for computationally efficient analysis of large-scale phenotyping data.

## 2 Materials and Methods

### 2.1 Clinical data extraction

The data used in this study represents the majority of the patient notes of Children’s Hospital of Philadelphia (CHOP). All medical documentation is performed using a unified Electronic Health Record (EHR) system, Epic (Verona, WI), for all care documentation (www.epic.com). The EHR system documents contacts with patients, including telephone call records, refills, visits for laboratory and imaging, hospital admissions, and clinic and emergency room visits. Each contact point is referred to as an encounter. Patient data from the production medical record is then merged into a separate reporting database provided by Epic (Clarity) as well as an internal database system, the Clinical Data Warehouse (CDW). For research purposes, a subset of this data is modeled within Arcus, an institutional informatics platform at Children’s Hospital of Philadelphia [34]. Within the Arcus library system, the Arcus Data Repository (ADR) de-identifies the clinical data allowing researchers to access and conduct studies on de-identified data, reducing administrative burden and lifting Institutional Review Board (IRB) oversight requirements.

### 2.2 Mapping to Human Phenotype Ontology terms

Many commercial NLP systems struggle with recognizing clinical text, and those trained on clinical data are usually costly, proprietary, and lack customizable features. Apache Clinical Text Analysis and Knowledge Extraction System (cTAKES), an open-source project, tries to fill this gap [17]. Although cTAKES provided an NLP solution specialized in health data, the analysis of large sets of clinical notes is time-and resource-intense. Accordingly, in its default configuration, this framework is not practical for institution-size data, and many institutions use cTAKES only on a small subset of their records [35, 36].

We have developed a pipeline that is capable of running cTAKES on millions of clinical notes in parallel, allowing for utilizing institution-size data at full capacity [37]. We applied cTAKES 4.0.0 [38] with the default clinical pipeline on all CHOP clinical notes with two modifications. First, we used a dictionary based on the Human Phenotype Ontology, HPO release version 2020-10-12, as included with the 2021AA release of Unified Medical Language System (UMLS) [39]. Secondly, we included the NegEx negation annotator in addition to the default machine learning negation classifier available in cTAKES [40]. NegEx is a regular expression algorithm that uses a list of patterns in an attempt to better capture negated terms in the encounter notes by filtering out sentences containing phrases that falsely appear to be negation terms. This process allowed us to map each encounter in the EMR to a set of HPO terms at a large scale.

### 2.3 Assessing HPO term frequencies corrected for frequencies of higher-level phenotypic concepts

We calculated the frequency of each phenotype by dividing the number of individuals mapped with that HPO term in at least one encounter by the total number of individuals. However, assessing the baseline frequency does not fully represent the complete frequency of each phenotypic term as higher-level terms are not typically included in encounters. Clinical descriptions tend to be annotated with the phenotypic term at the greatest applicable level of detail. Consequently, the analysis of the direct translation of clinical records into HPO terms underestimates the frequency of higher-level, conceptually broader, phenotype terms. However, it can be reasoned that these higher-level terms are often essential as they may capture large groups of individuals with phenotypically broad but clinically or biologically important similarities. Accordingly, we performed a method referred to as “propagation” that we have used extensively in past work [6, 25, 41-45]. In brief, for each individual’s set of explicitly annotated HPO terms, we add all HPO terms that can be inferred as applicable by taking the union of all those terms encountered following all possible paths along “is a” relationships to the root of the HPO. For example, if an individual was coded with *Mild global developmental delay (HP:0011342)*, this technique ensures counting this individual when calculating the frequency of all higher-level terms, including *Global developmental delay (HP:0001263)* and *Neurodevelopmental delay (HP:0012758)*. The term frequency of the propagated terms provides a more accurate representation of the true term frequencies for concepts coded in the HPO [44]. We refer to the frequency derived from the propagated counts as propagated frequency. Thus, the propagated frequency of the root term in the HPO, *freq* (*HP*: 0000001), is always equal to 1.

### 2.4 Similarity analysis using the Resnik Score

Various frameworks have been introduced to automatically measure the semantic similarity between concepts [46, 47]. Among them, many focus on deriving statistical information from cohorts and combining them with lexical resources and knowledge graphs [48]. These techniques have been effective in NLP tasks such as information extraction based on WordNet [49]. In 1995, Resnik introduced information content (IC) as a measure of specificity for a concept [22]. The IC of each concept represented in an ontology can be calculated based on the occurrence frequency of that concept in a large and relevant corpus. As a result, a generic concept would be associated with a lower IC value, and a very specific and rarely encountered concept would have a high IC. In calculating the IC of a concept, the frequency of all concepts encountered following all possible paths along “is a” hierarchy to the main concept should be counted. Formally, the IC of a concept can be calculated as:

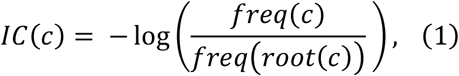

where *c* represents the concept, *root* represents the root of the hierarchy, and *freq*(.) returns the frequency value of a given concept.

Having the IC, the Resnik score computes the similarity between concepts *c*1 and *c*2 based on their Most Informative Common Ancestor (MICA), the concept that subsumes *c*1 and *c*2 and has the maximum IC. Formally,

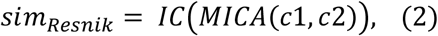

where *MICA*(*c*1, *c*2) is the lowest common subsumer of concepts *c*1 and *c*2, and *IC*(.) returns the information content of a concept.

In the HPO, represented as a DAG, phenotypes closer to the root are more general and have a lower IC value. The *MICA* of two phenotypes is the least common ancestor (LCA) of those phenotypes in the HPO. As a result, the quantity of shared information between two phenotypes would be higher if the LCA is a rare term and has a high IC. In some cases, the Resnik score cannot make a fine-grained distinction as many phenotypes may share the same LCA and thus receive equal similarity scores. In addition, the frequency values involved in calculating the information content may suffer from different sources of biases in the reference cohort, including but not limited to selection bias, information bias, and confounding bias [50]. Although some bias sources can be identified, and their potential impact can be assessed, they may require extensive data cleaning and supervised analysis, which are very costly for institutional data. Additionally, many sources of bias are hidden. Thus, techniques like Resnik that directly use the phenotypic frequencies in measuring similarities are susceptible to such biases.

### 2.5 Node embeddings using Node2Vec

Inspired by advancements in word embeddings for NLP tasks, the Node2Vec model maps nodes in the graph to a d-dimensional vector space [30]. There are several sampling strategies that are used to explore nodes in the graph [30, 51, 52]. Different sampling strategies generate different sentence-like samples that result in distinct learned representations. Some of these strategies rely on a rigid notion of a network neighborhood and are insensitive to connectivity patterns unique to networks [51, 52]. As a result, they have shortcomings in generalizing different prediction tasks and graph structures. Therefore, it is essential to allow for a flexible sampling strategy to explore the graph and learn node representations with two rules in mind: (1) learning representations where nodes with similar roles in the graph receive similar embeddings, and (2) learning representations where nodes from the same substructure or community are set closer together in the space. Node2Vec introduces a biased randomized procedure that samples neighborhoods for each given node where the transition probability to the next node depends on both the current and previous node. This procedure overcomes the mentioned shortcomings and follows the listed rules. The generated biased random walks from each node preserve the mutual proximity among the nodes, effectively exploring a node’s local neighborhood. By running multiple biased random walks on each node, Node2Vec generates sets of sentence-like sequences, which serve as inputs to the Skip-gram model used in word embedding [29] (**Supplementary Method S1**).

The Skip-gram model aims to learn a continuous representation of words by optimizing a neighborhood likelihood objective in a semi-supervised fashion. The Skip-gram objective is based on a distribution hypothesis stating that words that appear in a similar context (windows over sentences) tend to have similar meanings. Particularly, similar words tend to appear in similar word neighborhoods and should be moved closer in vector space. For example, the words “epilepsy” and “illness” will likely be much closer in vector space than “table”, as they are much more likely to appear within the same windows of text. The biased random walks introduced in Node2Vec sample the nodes’ neighborhoods and serve the same purpose as context/sentences for the Skip-gram model.

### 2.6 HPO2Vec and Node2Vec+ as extensions of the Node2Vec framework

Phenotype embedding for HPO data has been developed by applying Node2Vec on the phenotypic nodes with weights equal to 1, a tool referred to as HPO2Vec [32]. Despite HPO2Vec’s promising results on specific phenotypes for a limited number of diseases, the equal weighting strategy prevented connection strength from being incorporated into exploring the graph, placing phenotypes equally close to their general and rare neighbors. A possible solution to this problem is to include connection strength in the form of edge weight in the HPO (see **Section 2.7**). While Node2Vec was designed to work on both weighted and unweighted (equal-weight) graphs, it does not distinguish weak connections from stronger ones and thus cannot detect cases where the potential next node is weakly connected to the previous one. The Node2Vec+ model proposed an extension of Node2Vec that resolved the issue with weak connections [53]. Intuitively, a connection is called “loose” based on some threshold edge value; however, it is hard to determine a reasonable threshold value for networks without having the distribution of edge weights of all nodes. Accordingly, they set a relative threshold for each node based on its surroundings. Formally, to evaluate if a connection (*u, v*) is loose, where *u* and *v* are two nodes in the graph, they compare its weight with the average edge weight from *u*, defined as:

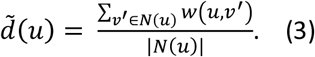

If 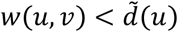, the edge, (*u, v*), is referred to as a loose connection. For simplicity, (*u, v*) is also considered a loose connection if (*u, v*) ∉ *E*. The extended bias factor is defined as:

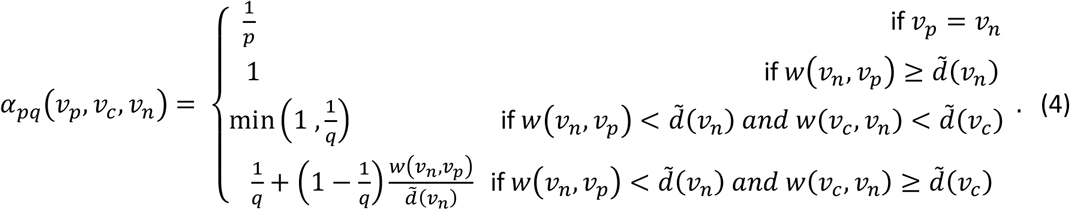

Note that for an unweighted graph, **Equation 4** acts as the bias factor in Node2Vec (**Supplementary Method S1)**, and there is no difference between the two methods. Also, similar to Node2Vec, if *p* and *q* are set to 1, the biased random walk will work as a simple first-order random walk.

### 2.7 Frequency-based human phenotype (FQ-HP) embedding

Here we propose a frequency-based human phenotype (FQ-HP) embedding model which consists of three steps, including: (1) updating the HPO graph by incorporating the propagated frequencies, (2) creating phenotype embeddings using Node2Vec+, and (3) calculating the similarity between phenotypes. Available studies on phenotype embedding, including HPO2Vec [32] and HPO2Vec+ [33] (see **Section 4**), apply their techniques to equally-weighted graphs which are equivalent to unweighted graphs. This results in choosing among surrounding phenotypes evenly rather than incorporating any connection strengths and priorities. This shortcoming would ultimately create equivalent similarity values between a phenotype with its rare child and the same phenotype with its relatively general neighbor (which could be another child or its parent). We call this category of techniques Equal-weight human phenotype (E-HP) embeddings and employ them in our evaluations. See **Section 3** for more details.

#### 2.7.1 Updating the HPO graph

The HPO version used in the current project contains 15,371 nodes (phenotypes) connected with unweighted and directed edges. To apply the Node2Vec+ algorithm, we created a copy of the HPO graph, including all 15,371 nodes and their accompanying 19,523 undirected edges. Assigning undirected edges helps the biased random walks in Node2Vec+ explore each given phenotype’s lower-level and higher-level neighborhoods. Additionally, we assigned weights to the edges of our graph. Weighted edges affect the probability of choosing the next nodes to visit in the biased-random walks. More specifically, at each step in the biased-random walk, if the algorithm needs to choose amongst a group of nodes to visit next, it will most likely choose the node (phenotype) with the strongest connection (largest weight) to the current node (**Equation S7**). We used the frequency calculated in **Section 2.3** to weight the edges. More precisely, the weight between two connected nodes is calculated by the minimum frequency value of the two nodes, formally defined as:

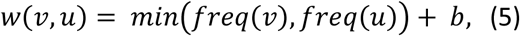

where *v* and *u* are two connected nodes (phenotypes), and *freq*(.) determines the frequency of a given phenotype. Here, *b* is a constant bias value that prevents zero weights. In our study, we used the difference between the maximum frequency and the second maximum frequency among all phenotypes as *b* (0.00146). The weighting mechanism introduced in **Equation 5** creates stronger connections between phenotypes with higher frequency values (more general terms) compared to rare terms. Furthermore, if phenotype *p* has multiple neighbors, its connection(s) to its parent(s) will be stronger than to its children — that are equally rare or rarer. Conceptually, this puts rare phenotypes further away in vector space compared to common ones as we argue that they represent unique information about a patient. We have provided the example of embedding the HPO terms *Generalized-onset seizure (HP:0002197)* and *Motor seizure (HP:0020219)*, which represent children of the phenotype *Seizure (HP:0001250)* in **Figure 1** and **Supplementary Method S1**.

**Figure 1.**
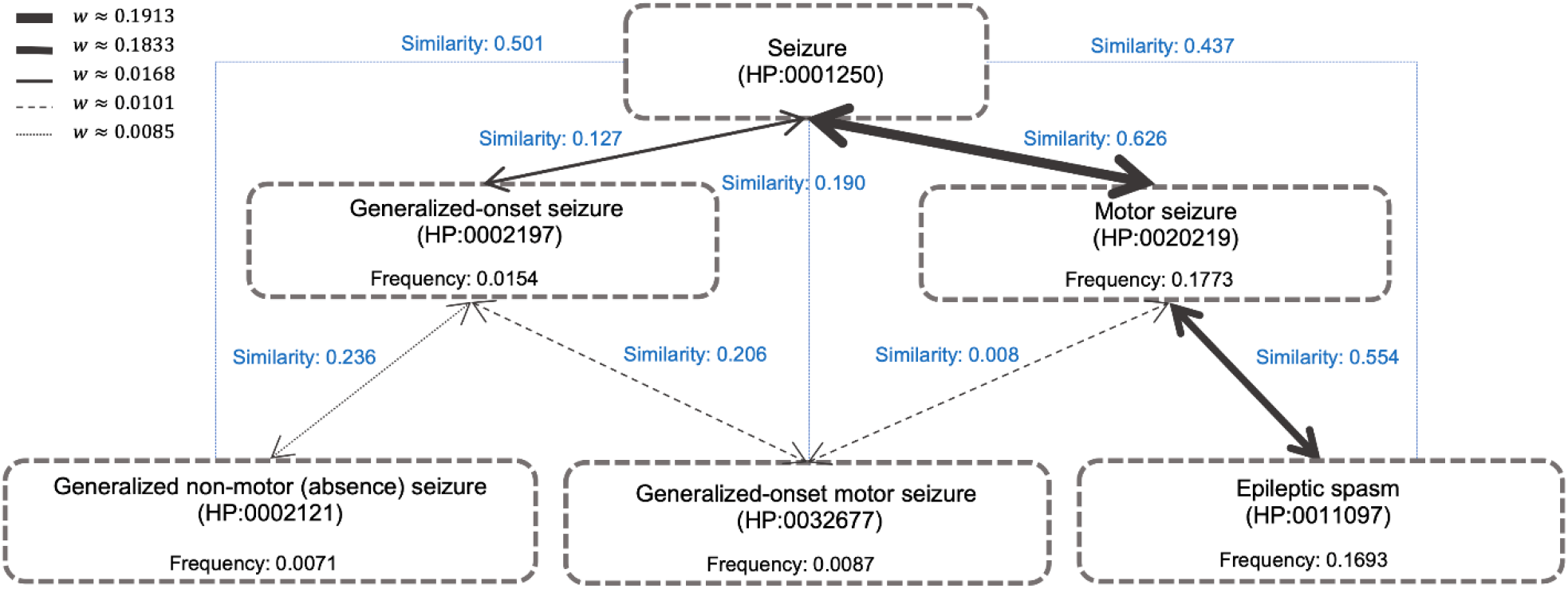
An illustration of the weighting mechanism on a portion of the HPO. The thickness of an edge represents its weight. Stronger weights have a higher chance of being selected by the biased random walks in the sampling algorithm. The propagated frequency values are calculated as described in **Section 2.3**. The cosine similarity values between each phenotypic pair are shown in blue. Higher similarity values represent a higher degree of closeness in the embedding space.

#### 2.7.2 Creating phenotype embedding

We used the Node2Vec algorithm, described in **Section 2.5**, with the extended bias factor introduced in Node2Vec+ (**Section 2.6**) to incorporate our weighting mechanism in embedding the phenotypes. We ran the embedding algorithm on all 15,371 phenotypes available in the HPO release 2020-10-12.

#### 2.7.3 Hyper-parameters

Since our task is unsupervised and there is no label involved in the fine-tuning process, we relied on the provided hyper-parameters used in HPO2Vec+ for the sampling strategy [33]. The select hyperparameters used in the sampling strategy as well as training the Skip-gram model are provided in **Table 1**.

**Table 1.** Select hyper-parameters used for the phenotype embedding methods.

| Hyper-parameter | Value |
| --- | --- |
| vector length (dimension) | 128 |
| $p$ | 1 |
| $q$ | 0.05 |
| number of walks | 10 |
| number of steps | 5 |
| learning rate | 0.001 |
| number of negative samples | 4 |
| batch size | 1024 |
| number of epochs | 15 |

#### 2.7.4 Representing the embedding space

With the embedding model, each phenotype can be mapped into the embedding space with its vector representation. However, the embedding space has high dimensionality, where dimensionality is defined by the hyper-parameter vector length. Thus, visualizing how phenotypes occupy the embedding space is impossible. We apply two dimensionality-reduction techniques, namely, Principal Component Analysis (PCA) [54] and t-distributed stochastic neighbor embedding (t-SNE) [55] to the embedding vectors to visualize them in 2-D and 3-D space. Since the dimensionality reduction techniques can potentially lose important information from the original data, we use them only for visualization purposes and work with the original vectors when comparing phenotypes and calculating their similarities.

#### 2.7.5 Calculating phenotype similarities

There are three main techniques available in the literature to calculate the similarity between two embedding vectors: Euclidean distance, Cosine similarity, and dot product. Among these techniques, the dot product is proportional to the vector length, which is a hyper-parameter of the sampling strategy. Thus, we only used the Euclidean distance and Cosine similarity to measure the similarities between the two vectors in our experiments (**Table 2**).

**Table 2.**
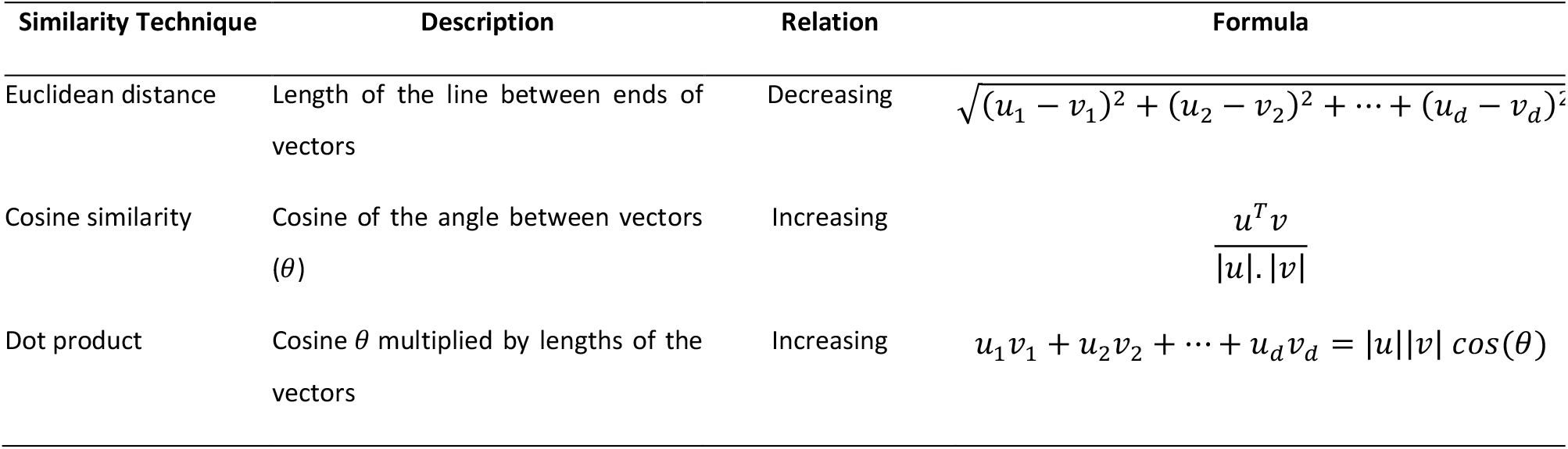
Techniques for measuring the similarity between two vectors *u* and *v* of size *d*.

## 3 Results

### 53 million patient notes were translated into 15,371 phenotypes

We analyzed data from 1,504,582 patients with a wide range of syndromes available on 53,955,360 electronic notes in the Arcus Data Repository version 1.4.4 [34]. By applying the cTAKES algorithm on all available full-text notes from the EMR, we extracted 8,425 distinct HPO terms with more than one occurrence and 9,477 distinct phenotypes with more than one propagated occurrence. Patient encounters over time and the histogram of the propagated frequency of phenotypes are presented in **Figure 2**.

**Figure 2.**
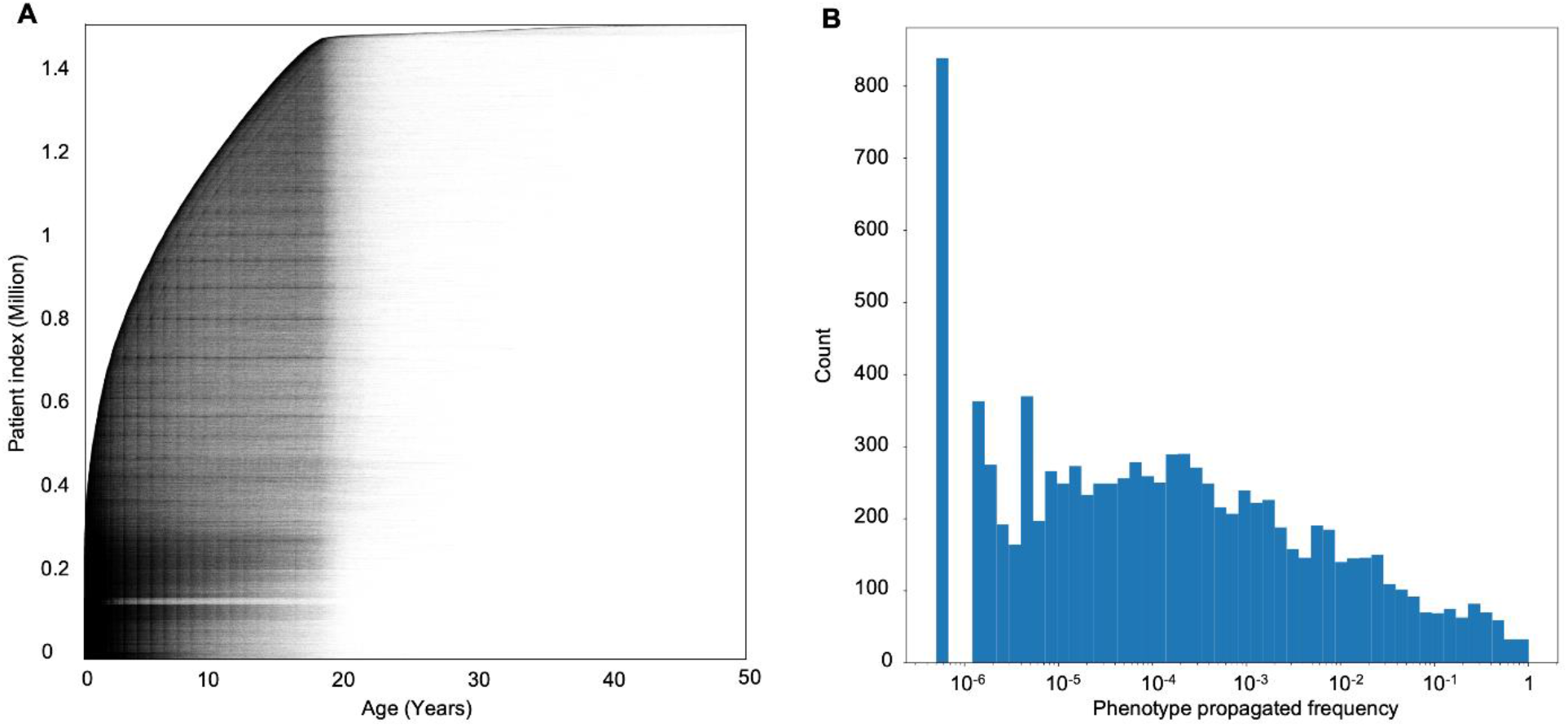
More than 53 million patient health records can be translated into phenotypes. **(A)** Individual data points in the Arcus Data Repository used for our analysis (n = 53,955,360). The X-axis shows patients’ age at the time of the encounter, and the Y-axis represents patients’ indices stacked and sorted by their age at the earliest encounter with a total of 1,504,582 individuals. **(B)** Histogram of the phenotypes’ propagated frequency available in the HPO, a total of 15,371 phenotypes. Note that the X-axis has a logarithmic scale, and the phenotypes with a frequency of 10^−6^ were not available in any of the encounters.

### Representing the Human Phenotype Ontology in a lower-dimensional space

We qualitatively evaluated our embedding method by visualizing the 15,371 phenotypes in the embedding space. Since we designed the embedding vectors to be of length 128, it is impossible to visualize the space directly. We used the PCA and t-SNE algorithms to reduce the embedding dimension to 2D and 3D. **Figure 3** displays a 3D representation of the original (128D) embedding space using PCA, where phenotypes that are closer together are more similar.

**Figure 3.**
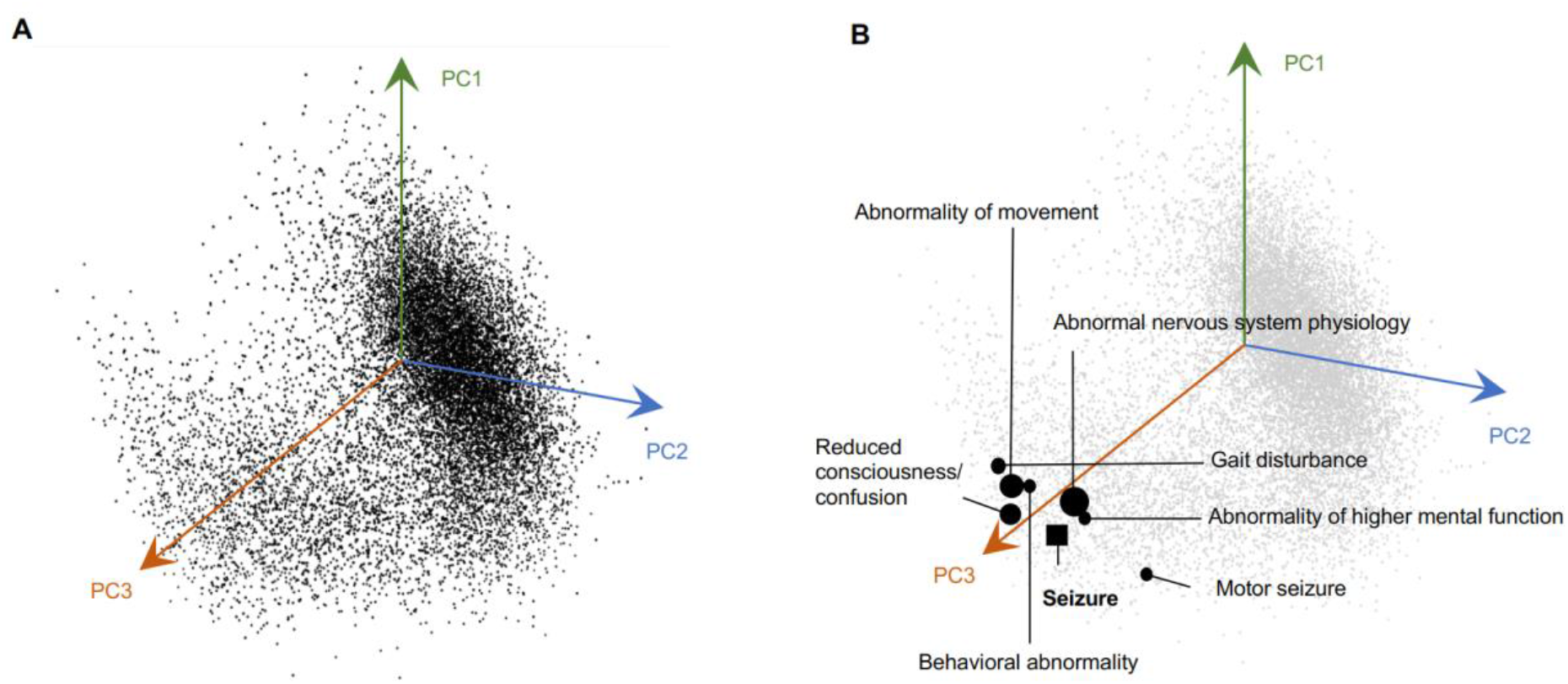
The HPO can be represented in a lower-dimensional space where similarities and differences between the phenotypes are preserved. **(A)** A 3D representation of all phenotypes in the embedding space using the PCA algorithm. The axes are based on the first three principal component vectors (PC) of the PCA algorithm. **(B)** Seven closest phenotypes to Seizure (HP:0001250) are marked in the 3D space. Closer phenotypes in the original space (128D) are represented with larger circles.

### Graph representation learning preserves relationships between phenotypes in the embedding space

We next assessed the Skip-gram objective to determine if proximity to similar contexts would drive the phenotypes to be closer in the embedding space. In doing so, we examined a group of phenotypes, including *Seizure (HP:0001250)* and its neighbors in the HPO, and measured their similarities in the embedding space. **Table 3** shows the similarity values between *Seizure* and some of its closest neighbors in the original embedding space. We calculated the similarity using the Cosine Similarity and Euclidean Distance metrics with larger values in the Cosine System representing a higher degree of similarity while smaller values in the Euclidean System representing shorter distances, thus greater similarities. Even in a lower-dimensional 3D space generated by PCA, the close neighbors of *Seizure* are evident (**Figure 3**). We also used the t-SNE algorithm to reduce the embedding space to three dimensions. Our results demonstrate that while t-SNE can preserve the similarity and difference between most of the phenotypes, tuning its hyper-parameters (perplexity, learning rate, and iteration number) to obtain a stable low-dimensional representation adds additional complexity that does not necessarily improve the representation (**Supplementary Figure S1**).

**Table 3.**
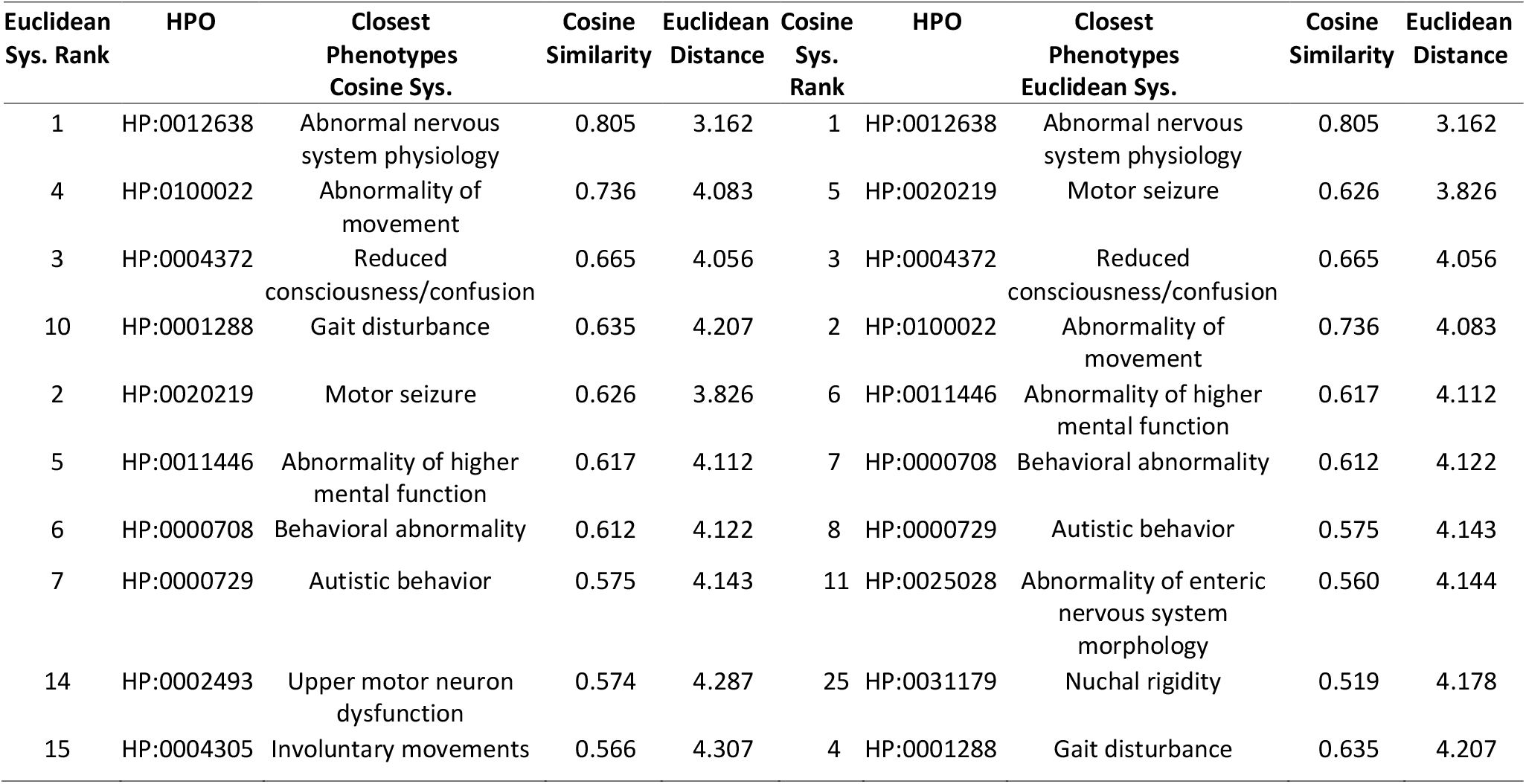
Closest phenotypes to the node Seizure (HP:0001250) in the embedding space using the Cosine Similarity System versus the Euclidean Distance System. While the majority of the phenotypes are common between the two systems, there are minor differences in their ranks.

| Euclidean Sys. Rank | HPO | Closest Phenotypes Cosine Sys. | Cosine Similarity | Euclidean Distance | Cosine Sys. Rank | HPO | Closest Phenotypes Euclidean Sys. | Cosine Similarity | Euclidean Distance |
| --- | --- | --- | --- | --- | --- | --- | --- | --- | --- |
| 1 | HP:0012638 | Abnormal nervous system physiology | 0.805 | 3.162 | 1 | HP:0012638 | Abnormal nervous system physiology | 0.805 | 3.162 |
| 4 | HP:0100022 | Abnormality of movement | 0.736 | 4.083 | 5 | HP:0020219 | Motor seizure | 0.626 | 3.826 |
| 3 | HP:0004372 | Reduced consciousness/confusion | 0.665 | 4.056 | 3 | HP:0004372 | Reduced consciousness/confusion | 0.665 | 4.056 |
| 10 | HP:0001288 | Gait disturbance | 0.635 | 4.207 | 2 | HP:0100022 | Abnormality of movement | 0.736 | 4.083 |
| 2 | HP:0020219 | Motor seizure | 0.626 | 3.826 | 6 | HP:0011446 | Abnormality of higher mental function | 0.617 | 4.112 |
| 5 | HP:0011446 | Abnormality of higher mental function | 0.617 | 4.112 | 7 | HP:0000708 | Behavioral abnormality | 0.612 | 4.122 |
| 6 | HP:0000708 | Behavioral abnormality | 0.612 | 4.122 | 8 | HP:0000729 | Autistic behavior | 0.575 | 4.143 |
| 7 | HP:0000729 | Autistic behavior | 0.575 | 4.143 | 11 | HP:0025028 | Abnormality of enteric nervous system morphology | 0.560 | 4.144 |
| 14 | HP:0002493 | Upper motor neuron dysfunction | 0.574 | 4.287 | 25 | HP:0031179 | Nuchal rigidity | 0.519 | 4.178 |
| 15 | HP:0004305 | Involuntary movements | 0.566 | 4.307 | 4 | HP:0001288 | Gait disturbance | 0.635 | 4.207 |

### Incorporating phenotype frequencies in the HPO graph transfers likelihood distribution to embeddings

We next analyzed the effects of incorporating weights in the HPO graph. In doing so, we first calculated pair-wise cosine similarity values between all phenotype pairs. We observed changes in the similarity values when using the FQ-HP embedding (our technique) compared to the E-HP embedding (the conventional equal weights). **Figure 4A** represents the changes in the cosine similarity values between a sample phenotype (*Infantile spasms HP:0012469*) and all other phenotypes in the HPO. In this example, the range of cosine similarities is wider for FQ-HP embedding (our technique, y-axis) compared to the E-HP embedding (the conventional equal weights, x-axis). Considering that cosine similarity is bound by a constrained range of -1 and 1, the increased range for FQ-HP (y-axis) suggests that more phenotypes have a notable, more extreme positioning with respect to the main phenotype in the embedding space. We observed various patterns of changes in the similarity values on different phenotypes (**Supplementary Figure S2**).

**Figure 4.**
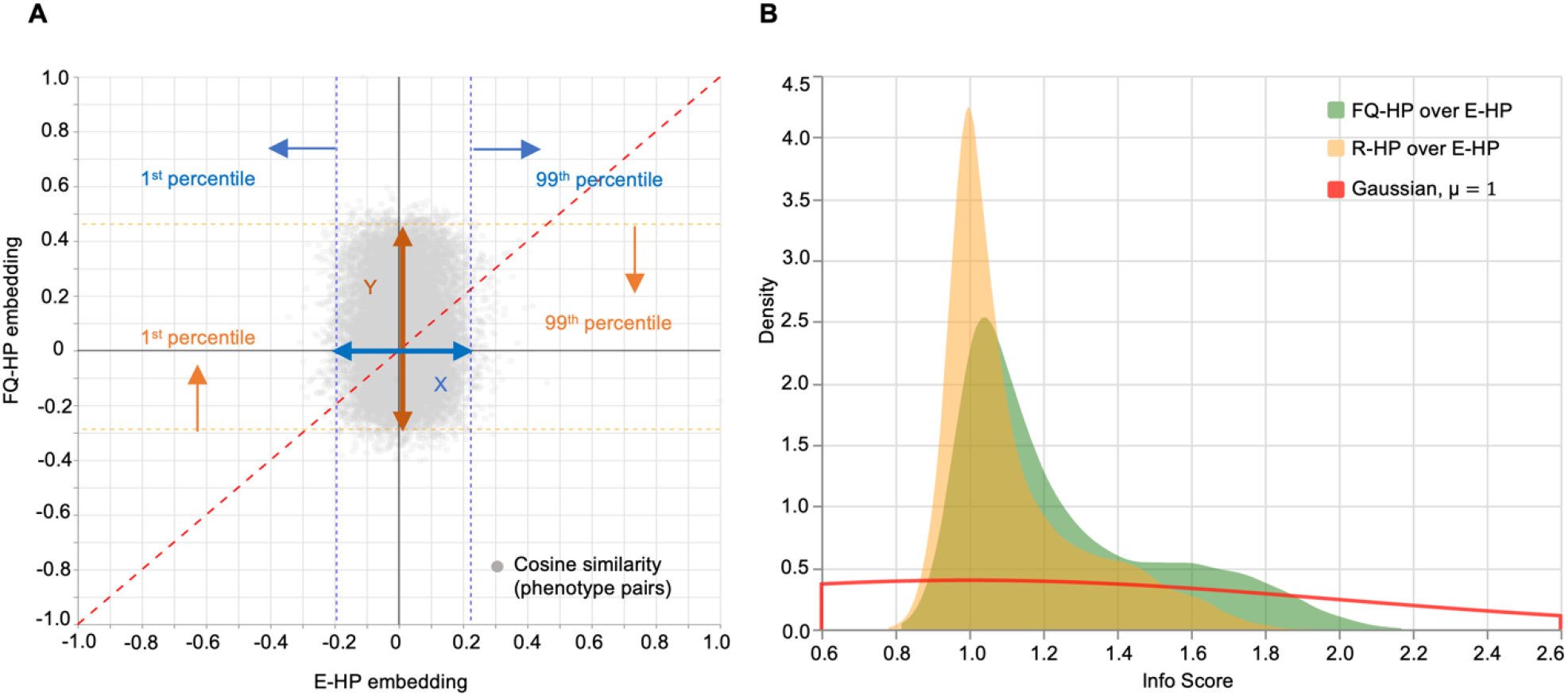
Incorporating phenotypes frequencies in the HPO graph transfers likelihood distribution to embeddings. **(A)** Pair-wise cosine similarity changes between a sample phenotype, *Infantile spasms (HP:0012469)*, and all other phenotypes in the HPO using the FQ-HP and E-HP embedding methods. *Info Score* captures the similarity changes between these two methods by dividing the similarity range using FQ-HP by the similarity range using E-HP, i.e., Y/X. **(B)** A comparison between the PDF of the *Info Score* of FQ-HP over E-HP (shown in green) and the *Info Score* of Random Gaussian weights (R-HP) over E-HP (shown in yellow). The PDF of *Info Score of* FQ-HP/E-HP with a smaller peak around 1 and a longer tail indicates that FQ-HP could generate reliable phenotype clusters in the embedding space and shows the importance of assigning weights that represent the actual distribution of the phenotypes.

In order to assess the patterns of distributions between FQ-HP and E-HP across all phenotypes, we defined a metric that compares the ratio between the range for 99% of all cosine similarity values for FQ-HP and E-HP, the ratio of Y to X as demonstrated in **Figure 4A**. We referred to this metric as “Info Score” (**Supplementary Method S3)** and compared the distribution of Info Scores of FQ-HP over E-HP with that of embeddings with randomly assigned weights (R-HP) over E-HP. The observed Info Score distribution of FQ-HP/E-HP (**Figure 4B, green**) suggests stronger similarity values compared to R-HP/E-HP (**Figure 4B, yellow**), indicated by the “longer tail” of the FQ-HP/E-HP distribution. This difference between embeddings using true frequencies and random frequencies indicates that, on average, the spectrum of similarities is broader when using frequency-based embedding. Accordingly, when using the additional information of term frequencies, the existing proximities and distances in the phenotypic graph can be represented more efficiently.

### Frequency-based phenotype embeddings improved recognition of expert-curated phenotype similarities

Next, we assessed how similarity using embeddings compared to human assessment of clinical similarities. To evaluate the utility of the frequency-based phenotype embeddings over conventional methods to assess clinical similarity, we generated an expert-curated dataset comparing a reference to two choices of phenotypes, here referred to as candidate phenotypes. The domain experts were asked to assess which of the two choices is more closely related to the reference phenotype in their opinion. They were also given the option to indicate uncertainty if they viewed the candidate phenotypes as equally related to the reference phenotype or found it impossible to choose. We reasoned that this would generate a gold standard dataset that provides a valuable framework to assess the utility of similarity-measuring algorithms. In brief, an algorithm that is more aligned with the expert-curated dataset is considered superior to an algorithm that results in less overlap.

To generate such a gold standard dataset, we provided a group of 13 domain experts in epilepsy and neurogenetics, including 12 clinicians and a researcher, with 100 neurology-related reference terms (i.e., *Abnormality of the nervous system (HP:0000707)* and all child terms). These terms were balanced between common and rare phenotypes as well as phenotypes above or below in the phenotypic hierarchy compared to the candidate phenotypes (see **Table 4** and **Supplementary Table S1**). In total, this resulted in 1,300 individual phenotype prioritizations. Across each phenotypic trio, a voting system was implemented, assigning the most frequent decision among the expert raters as the gold standard.

**Table 4.** Scenarios for generating expert-curated phenotypic trios.

| Scenario | Abbreviation | Description | #{total:100} |
| --- | --- | --- | --- |
| S1 | $CR_D R_D$ | Common<sup>1</sup> reference with two rare candidates and both candidates as descendants | 10 |
| S2 | $CR_D R_{\sim D}$ | Common reference with two rare candidates and only one candidate as a descendant<br>(positive control) | 5 |
| S3 | $CR_{\sim D} R_{\sim D}$ | Common reference with two rare candidates and neither as a descendant | 5 |
| S4 | $CC_D C_D$ | Common reference with one rare and one common candidate and both candidates as descendants | 5 |
| S5 | $CC_{\sim D} C_{\sim D}$ | Common reference with two common candidates not descending from the reference | 5 |
| S6 | $R_{\sim A \sim A} R_{\sim A \sim A} R_{\sim A \sim A}$ | Rare reference with two rare candidates, none being an ancestor of either of the other two terms | 50 |
| S7 | $RC_A C_{AA}$ | Rare reference with two common candidates, both being ancestors and one candidate being an ancestor of the other (positive control) | 5 |
| S8 | $RC_A C_{\sim A}$ | Rare reference with two common candidates, one being its ancestor and the other not (positive control) | 5 |
| S9 | $RC_A R$ | Rare reference with one common candidate and one rare candidate with the common candidate being an ancestor of the reference (positive control) | 5 |
| S10 | $RC_{\sim A \sim D} R_{\sim A \sim D}$ | Rare reference with one common candidate and one rare candidate and both unrelated | 5 |

<sup>1</sup> Common indicates phenotypes with the relative propagated frequency $> 20\%$ with respect to Abnormality of the nervous system (HP:0000707).

Next, we performed a leave-one-out analysis [56] and identified a 59.85% agreement between our raters. This value provided a reference for the expected accuracy of similarity-measuring algorithms. While this reflects more than 1.5 × accuracy of random agreement (38.46%), it still indicates a wide variability in the clinical assessment by expert reviewers (**Supplementary Method S4**).

We then assessed the similarity-measuring algorithms in various scenarios, comparing the conventional Resnik algorithm, phenotype-embedding using equal weight (E-HP), and frequency-based phenotype-embedding (FQ-HP). For the E-HP and FQ-HP algorithms, we also included a variation with a safety threshold (ST) that would declare ties if the distances between the two candidates and the reference phenotype were close (<0.06 in the cosine system). This safety threshold was implemented to prevent false-positive prioritizations based on marginal frequency differences of the candidate phenotypes and was chosen using a trial-and-error strategy.

We found that FQ-HP, with an overall accuracy of 67.8%, surpasses other similarity-measuring algorithms in matching the experts’ gold standard (**Figure 5A**) and is even higher than the agreement level between our raters. When applying the safety threshold, we see a significant difference between the performance of FQ-HP(ST) and E-HP(ST) with an accuracy of 62.4% and 47.4%, respectively. This result emphasizes the potential of incorporating frequencies in phenotype embeddings.

**Figure 5:**
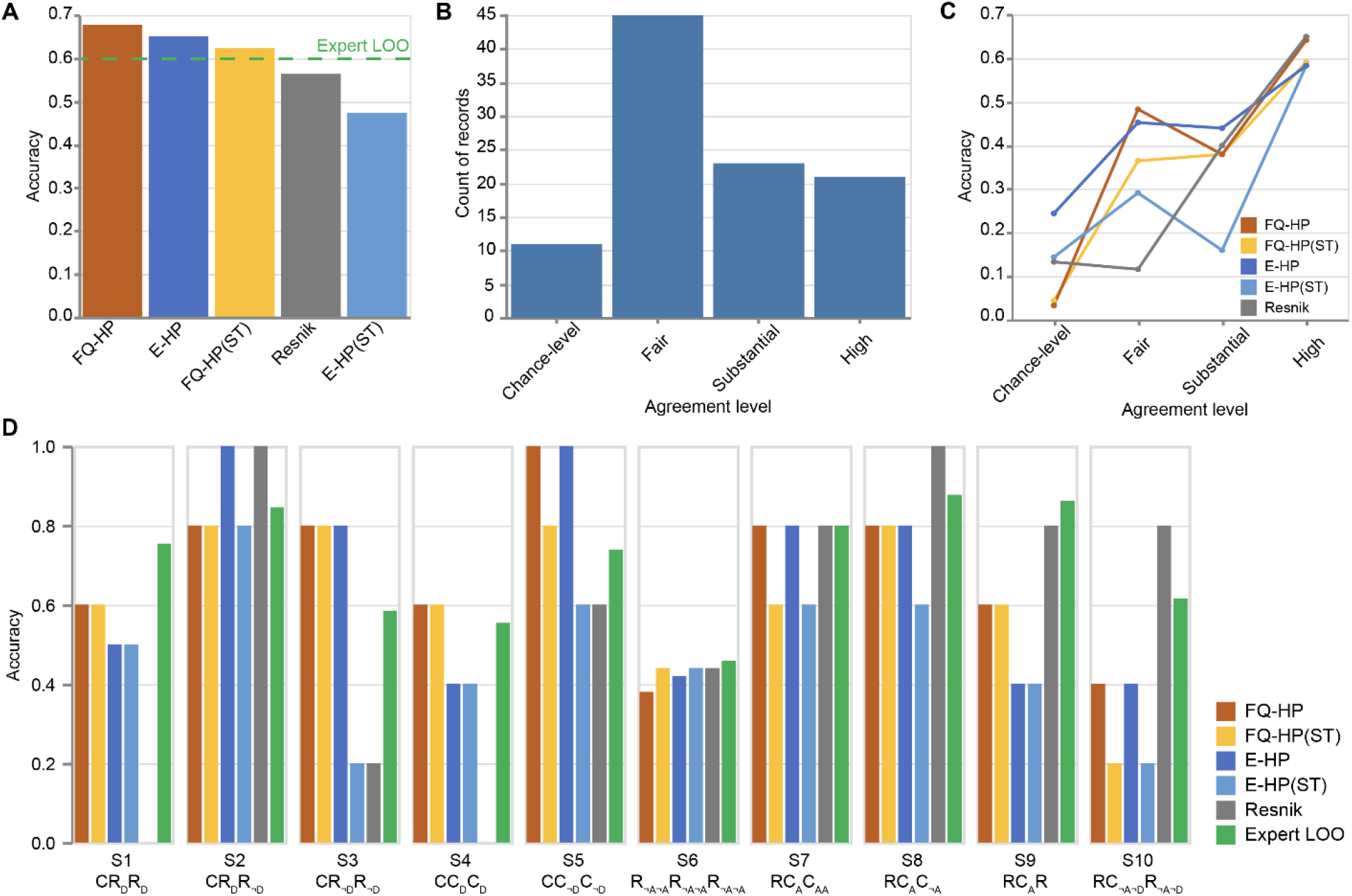
FQ-HP embeddings improve recognition of expert-curated phenotype similarities. **(A)** A comparison on the overall accuracy of the similarity-measuring algorithms on 100 expert-curated phenotypic trios. The accuracy of our method (FQ-HP) surpasses the other similarity-measuring techniques where black bars show 95% confidence intervals, and the green dashed line represents the experts’ agreement level. **(B)** Histogram of experts’ agreement level for the 100 trios. Almost half of the records belong to the fair-level agreement category, indicating a wide variability in the clinical assessment of the experts and the challenging nature of such evaluations. **(C)** Performance of the similarity-measuring algorithms on the records of each agreement level. FQ-HP and Resnik top the other techniques in substantial and high agreement levels. Also, FQ-HP achieves better accuracy than other techniques in the fair-level agreement. **(D)** Comparison on the scenario-based accuracy of the similarity-measuring algorithms. The conventional Resnik method performs better than the embedding algorithms in three scenarios while failing on four other scenarios, two of which with 0% accuracy (**Table 4**).

Next, we categorized our 13 experts’ decisions into four categories based on their agreement level (**Figure 5B, Supplementary Method S4**). We examined the performance of the similarity-measuring algorithms on the records of each agreement level. In short, if a similarity-measuring method performs inaccurately even when the experts have a high-level agreement, it would be considered a low-quality method. We found FQ-HP and Resnik to be more accurate than the other techniques in the “substantial” and “high” agreement levels. Additionally, FQ-HP achieved better accuracy than other techniques in the “fair-level” agreement. Since the “fair-level” is the most common category based on our human experts, a similarity-measuring technique needs to perform at least at this level on these challenging records (**Figure 5C**).

Finally, we also analyzed the efficacy of similarity-measuring algorithms in specific scenarios that were used in generating the expert-curated phenotypic trios (e.g., the combination of common and rare term frequencies as well as term hierarchies; **Table 4**). We found that the conventional Resnik method performed best for scenarios when only one candidate was in a hierarchical relationship with the reference term, positive controls provided in **Table 4**, providing accuracies close to expert assessments (S2^2^, S7^1^, S8, S9). However, for scenarios when the reference term is not in a hierarchical relationship with the candidates (S1, S3, S4, S5), embedding methods are superior. These findings indicate that the accuracy of various strategies to assess phenotypic relatedness may depend on the relationship and frequencies of the specific terms.

## 4 Discussion

This study aimed to assess the properties of phenotype embedding techniques to analyze the Human Phenotype Ontology (HPO) terms more computationally efficient and accurately reflect their complex biological and clinical relationships. We examined representations of HPO terms using node embeddings and compared the performance of embeddings with and without including HPO term frequencies derived from more than 53 million patient notes. Finally, we assessed the degree to which phenotype embedding methods align with expert opinion and how these methods performed in comparison to conventional phenotype similarity-measuring techniques. We demonstrate that incorporating phenotype frequencies from a large patient corpus resulted in phenotype embeddings surpassing conventional techniques in assessing phenotypic similarity and aligning more closely with assessments by domain experts. In contrast to conventional methods such as Resnik, our proposed method is fast and computation-efficient and can also be used in many downstream tasks such as patient similarities without any recalculation for new patients, directly from the embedding space. In summary, our results demonstrate that incorporating phenotypes frequencies from a large patient corpus as a weighting mechanism can transfer the phenotype distributions to the embedding space and ultimately provide a superior representation that aligns more closely with domain experts.

Our study has four main findings. First, we used Human Phenotype Ontology (HPO) to analyze clinical data in our study, which has its own advantages and disadvantages. HPO contains more than 15,000 clinical phenotypic terms with defined semantic relationships, developed to standardize their representation for phenotypic analysis. In addition, HPO is modeled as a directed acyclic graph (DAG) in which each phenotype is presented as a node and is connected to its parents by “is a” relationships using directed edges. Despite the fact that this structure facilitates phenotypic analysis, HPO is limited to symptoms, and diseases and syndromes are not directly mapped to its structure. While this limitation affects the in-depth analysis of clinical features, incorporating more than 15,000 phenotypic terms in our analysis allowed us to cover a wide range of complex clinical relationships.

Second, we demonstrate that clinical information from full-text patient notes can be translated at scale through Natural Language Processing. In our study, we mapped the entirety of the available full-text patient notes of a large, tertiary pediatric care network to Human Phenotype Ontology terms, generating the most comprehensive estimates for the frequencies of clinical concepts in the pediatric population currently available. Given the increasing use of HPO for both common and rare diseases, having valid term frequencies will be essential for the development of algorithms aimed to assess phenotypic overlaps, such as the assessment of autistic-like traits [57] and rare neurodevelopmental disorders [58].

Third, we demonstrate that phenotype embedding is a useful tool to represent the >15,000 HPO concepts in a lower-dimensional space while preserving existing phenotypic relationships. Additionally, the inclusion of observed term frequencies improves the vector embedding. This was demonstrated by the embedding’s ability to generate meaningful proximities and distances between clinical terms. When compared to randomly assigned term frequencies, the inclusion of term frequencies in the embedding allows for more closely related phenotypes to be “drawn in” and more distantly related phenotypes to be “pushed out.” This exemplifies how additional information incorporated in the embedding technique such as term frequency can provide more specificity and nuance in the connections between phenotypes. Future iterations including temporal components [11] or inclusion of additional data such as medication and procedure information may further improve phenotype embeddings and exceed the currently available frameworks.

Fourth, we demonstrate that the phenotype embedding methodology is largely superior to conventional similarity-measuring techniques; however, its strengths and weaknesses are dependent on the specific clinical scenario. For all comparisons, frequency-based embedding is slightly superior to equal-weight embeddings and Resnik similarity. However, if the reference terms are in a hierarchical relationship with one of the candidates, all embedding techniques are on par with or inferior to Resnik. We reason that this strong effect of specific scenarios reflects the inherent mechanics of conventional methods. For example, the Resnik algorithm cannot generate meaningful similarities in situations when terms are not in a direct hierarchy, which is demonstrated by our results for Scenarios S1 and S3-S5. In contrast, frequency-based embeddings are susceptible to potential errors in frequency assessments, which may be critical when rare references are used.

Despite the large amount of data, our mapping algorithm and proposed similarity measuring approach have several limitations. First, we observed several phenotypes with unexpectedly large frequency values in our cohort; for example, *Seizure (HP:0001250)* had a propagated frequency of 0.4248. Given the large and diverse source of encounter notes available in our corpus and the multi-layer phenotype extraction process, we could not trace back these instances and update the extracted phenotypes in the encounters and their consequent frequencies. These frequencies could potentially shift the biased random walks, reducing their effectiveness in exploring neighborhoods; however, our proposed embedding method attempts to reduce the effect of such potential artifacts in the data by assigning the minimum propagated frequency of two phenotypes as their edge weight as well as using relative weights in graph exploration.

Furthermore, like other distance-based embedding algorithms, the biased random walks in our method are bound within a defined walk length and tend to exploit the local neighborhood more than exploring further nodes which results in treating the denser sub-graphs of HPO differently. Potential future solutions to overcome this problem may involve flexible exploring criteria, using dynamic *p* and *q* hyper-parameters depending on the sub-graph structure to navigate both sub-structures of the HPO more evenly. Unlike conventional machine learning problems, we did not have ground-truth similarity values for the phenotype pairs to tune the hyper-parameters used in our model, a limitation imposed by the structure of the data assessed in this study. These limitations shed light on the importance of incorporating domain experts’ knowledge in tuning the model parameters and potentially achieving higher accuracy.

Moreover, in evaluating our proposed method against domain experts’ judgments, we limited ourselves to neurological phenotypes. Given the expertise of our domain experts, we only validated the similarity-measuring techniques for phenotypes under “*Abnormality of the nervous system (HP:0000707)*” with thirteen domain experts in epilepsy and neurogenetics. The agreement level among our domain experts suggests that measuring phenotypic similarity could be a challenging task even for experts in a particular field. This is particularly the case when experts are asked to compare pairs of phenotypes that are distant and hard to compare on anatomic or functional grounds. Having access to a larger and more diverse pool of domain experts could help us evaluate our model more thoroughly and obtain insights into overcoming such challenges. Additionally, based on the unsupervised nature of our embedding algorithm, we expect our proposed technique to work with approximately the same efficacy on phenotypes related to other fields of medicine.

In summary, we demonstrate that assessing clinical similarity in large EHR-derived datasets using phenotype embeddings may have specific advantages in scenarios where other similarity-measuring techniques have difficulties. We believe incorporating information from patient records, such as the frequency of co-occurrence of phenotypes, will improve phenotypic similarity-measuring techniques. Incorporating additional disease-phenotype information has been explored before in the form of adding additional edges to the HPO in embedding phenotypes, a method referred to as HPO2Vec+ [33]. Although this technique was found to be more successful than embedding phenotypes purely based on the HPO, it required expert-curated data, has been evaluated on a limited set of phenotypes and diseases, and is not practical for hospital-size datasets. Our goal, however, is to provide unsupervised methods that are more robust and generalizable to large-scale data. Given the increasing availability of electronic health records and phenotype extracting techniques, our phenotype embedding algorithm has the potential to be used in downstream tasks such as accessing patients’ similarities, identifying novel genetic etiologies, and ultimately providing better care to individuals.

## 5 Acknowledgments and Funding

I.H. was supported by The Hartwell Foundation (Individual Biomedical Research Award), the National Institute for Neurological Disorders and Stroke (K02 NS112600, U24 NS120854, U54 NS108874), the Intellectual and Developmental Disabilities Research Center (IDDRC) at Children’s Hospital of Philadelphia and the University of Pennsylvania (U54 HD086984), and by the German Research Foundation (HE5415/3-1, HE5415/5-1, HE5415/6-1, HE5415/7-1). Research reported in this publication was also supported by the National Center for Advancing Translational Sciences of the National Institutes of Health (UL1 TR001878), by the Institute for Translational Medicine and Therapeutics’ (ITMAT) at the Perelman School of Medicine of the University of Pennsylvania, and by Children’s Hospital of Philadelphia through the Epilepsy NeuroGenetics Initiative (ENGIN). This research was funded in whole, or in part, by the Wellcome Trust [203914/Z/16/Z] supporting D.L.S.. For the purpose of Open Access, the author has applied a CC BY public copyright license to any Author Accepted Manuscript version arising from this submission.

## 1 Supplementary Method S1

### 1.1 Skip-gram

Formally, given a graph *G* = (*V, E*) where *V* is a set of vertices (also known as nodes), and *E* is a set of paired vertices (also known as edges), let *f* be the mapping between each node *v* ∈ *V* to its feature representation of size *d* where *f*: *V* → ℝ^*d*^. Here, *d* is a parameter that defines the dimension of the feature representation. For each node *v* ∈ *V, N*_*s*(*v*)_ ⊂ *V* is a neighborhood with sampling strategy *S*, and the objective function maximizes the log-probability of reconstructing this neighborhood. Formally defined as:

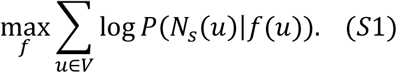

Two assumptions are made to solve the optimization problem presented in **Equation S1** in polynomial time. First, the likelihood of observing a neighborhood node is independent of observing any other neighborhood node given the source node’s feature representation, as:

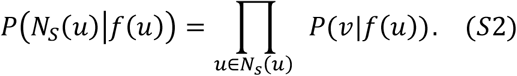

Second, a source node and neighbor node have a symmetric effect over each other; thus, their probabilities can be parameterized as the Softmax normalized inner product and can be written as:

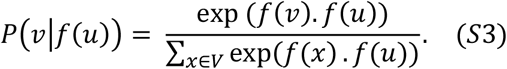

The objective function presented in **Equation S1** can be simplified with the conditional independence assumption and parameterization of the probabilities as introduced in [1] as follows:

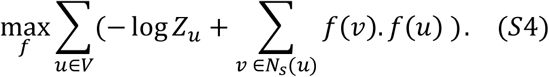

Calculating the partition function for each node, *Z*_*u*_ = ∑_*x*∈*V*_ *exp*(*f*(*x*). *f*(*u*)), is not practical, especially for larger networks.

In Node2Vec, they approximate the objective function using the negative sampling algorithm introduced in [2]. Instead of calculating the probability of co-occurrence of all node pairs, the negative sampling algorithm attempts to increase the co-occurrence probability of the sample node with its neighbors and decrease that probability with *k* randomly selected nodes from the graph. The simplified objective function presented in **Equation S4** can then find the d-dimensional representation of each node by employing the sampling strategy *S* and running Skip-gram with negative sampling.

Although Skip-gram revolutionized applications of NLP, it has two main drawbacks. (1) Skip-gram cannot capture polysemy, a word with multiple meanings, since it represents each word as a single vector. (2) It fails to identify compound word phrases. For example, the word “ice cream” should have a different representation than the words “ice” and “cream.” As we use the HPO terms as a representation of phenotypic terms in our phenotype embedding method, we do not face the aforementioned drawbacks.

### 1.2 Sampling strategy

The Skip-gram model was developed for inputs of text, where a neighborhood of a word is a sliding window on its surrounding words in unstructured text. In order to make graph structures amenable to the Skip-gram model, Node2Vec introduces a biased randomized procedure that samples neighborhoods for each given node. Unlike previous studies such as [3], where the transition probability to the next node, *v*_*n*_, only depends on the current node, *v*_*c*_, i.e., *P*(*v*_*n*_|*v*_*c*_), in a biased random walk the transition probability depends on both the current and the previous node, *v*_*p*_, i.e., *P*(*v*_*n*_|*v*_*c*_, *v*_*p*_), formally defined as:

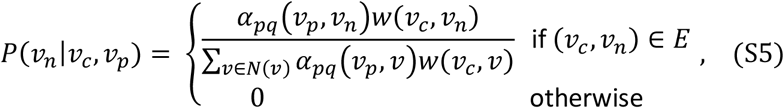

where *w*(*u, v*) represents the edge weight between nodes *u* and *v, N*(*v*) represents the sampled neighborhood for node *v*, and the bias factor *α*_*pq*_ is defined as:

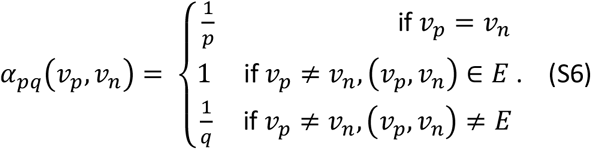

Parameter p controls the likelihood of immediately revisiting a node in a biased random walk, where larger values, >*max*(*q*, 1), encourage exploration in the graph. On the other hand, parameter q controls the exploitation criteria where larger values, >1, bias the random walk towards nodes that are closer to the previous node, *v*_*p*_.

In the case of a weak connection between nodes *v*_*p*_ and *v*_*n*_, i.e., 0 < *w*(*v*_*p*_, *v*_*n*_) ≤ 1, Node2Vec considers it a normal connection with a bias factor of 1 rather than 1/*q*. This case shows that Node2Vec does not discriminate weak connections from stronger ones and could not detect cases where the potential next node has a loose connection with the previous node.

## 2 Supplementary Method S2

### 2.1 Example of HPO embedding of phenotypic terms

*Generalized-onset seizure (HP:0002197)* and *Motor seizure (HP:0020219)* are represented as the children of *Seizure (HP:0001250)* in the HPO with a cumulative frequency of 0.0154 and 0.1773, respectively, in our corpus. These frequencies imply a stronger connection between *Seizure (HP:0001250)* and *Motor seizure (HP:0020219)* than Seizure (HP:0001250) and *Generalized-onset seizure (HP:0002197)*, which could be captured by our weighting mechanism.

Furthermore, *Generalized-onset seizure (HP:0002197)* has two children, *Generalized non-motor (absence) seizure (HP:0002121)*, with a frequency of 0.0071, and *Generalized-onset motor seizure (HP:0032677)*, with a frequency of 0.0087, which is also a child of *Motor seizure (HP:0020219). Epileptic spasm (HP:0011097)* is another child of *Motor seizure (HP:0020219)* with the frequency of 0.1693, which indicates a more robust representation of this phenotype in our corpus.

Based on the frequency of these nodes, *Motor seizure (HP:0020219)*’s connection is much stronger to its parent, *Seizure (HP:0001250)*, implying generalization in phenotypic concepts, with *w* = 0.626, than to either of its children, implying specialization in phenotypic concepts, with *w* equal to 0.008 and 0.554 (**Figure 1**).

These weights suggest placing *Motor seizure (HP:0020219)* and *Seizure (HP:0001250)* closer in the embedding space, resulting in a higher similarity value than *Motor seizure (HP:0020219)* to either of its children, *Epileptic spasm (HP:0011097)* or *Generalized-onset motor seizure (HP:0002197)*. The similarity values align with these suggestions and are marked in blue in **Figure 1**.

## 3 Supplementary Figure S1

**Figure S1.**
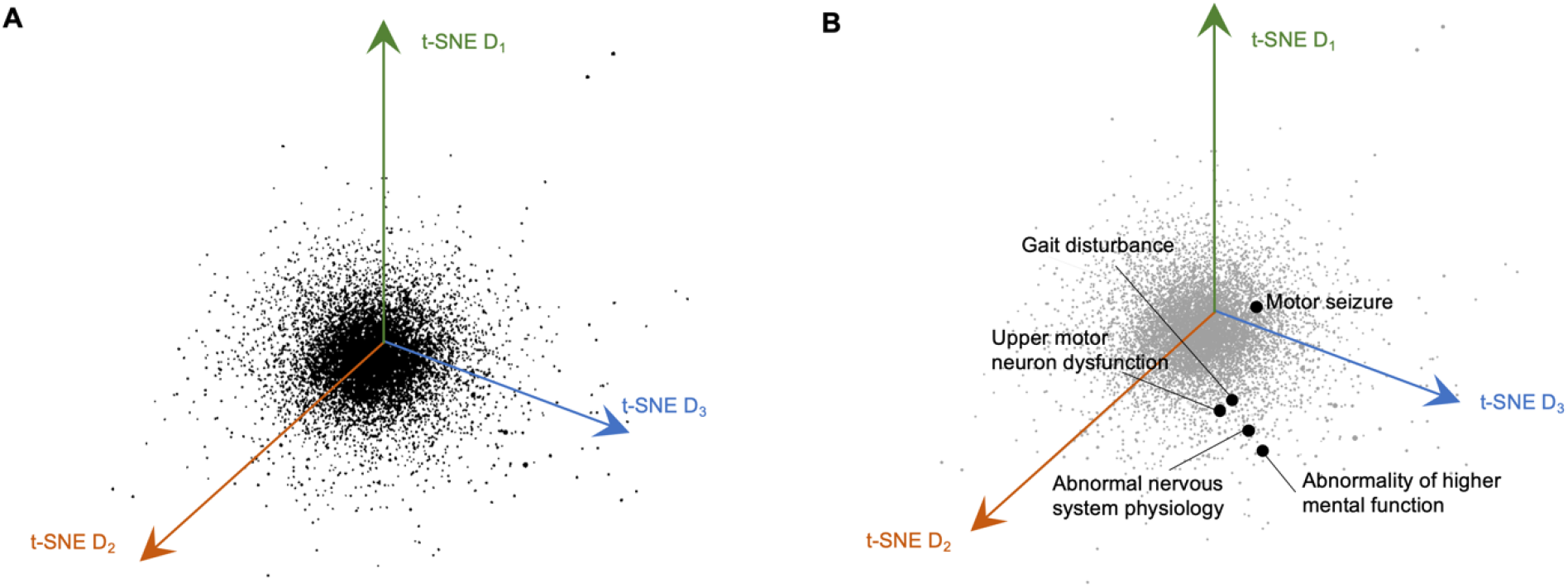
The HPO can be represented in a lower-dimensional space using the t-SNE algorithm, where similarities and differences between the phenotypes are preserved. **(A)** A 3D representation of all phenotypes in the embedding space using the t-SNE algorithm with 32 iterations. The axes are based on the first three t-SNE dimensions. We can see a change in the 3D representation compared to PCA 3D space (**Figure 3**). **(B)** The five closest phenotypes to *Seizure (HP:0001250)* are marked in the 3D space. While the closest phenotypes in PCA 3D space and t-SNE 3D space match, t-SNE requires hyper-parameter tuning, making it more costly compared to PCA.

## 4 Supplementary Figure S2

**Figure S2.**
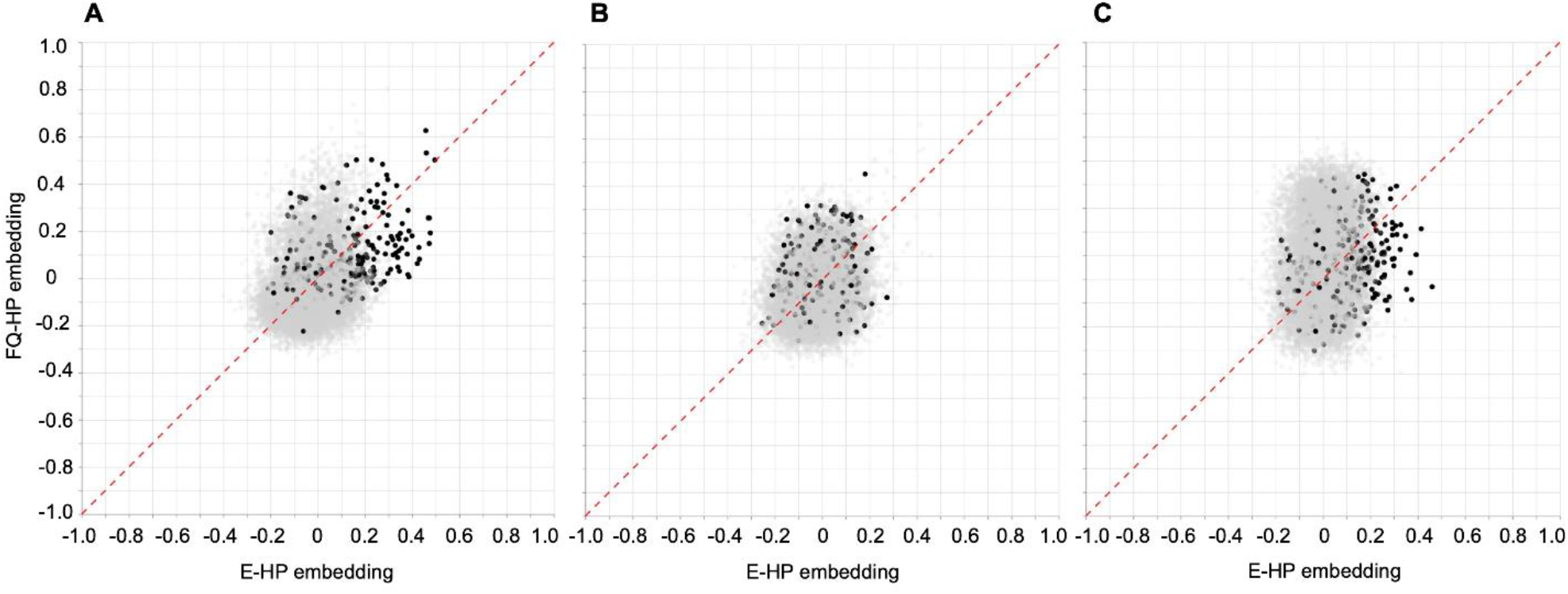
Different patterns of the cosine similarity value change using the E-HP and FQ-HP embedding. Pair-wise cosine similarity changes between a reference phenotype, **(A)** *Seizure (HP:0001250)*, **(B)** *Neurodevelopmental abnormality (HP:0012759)*, and **(C)** *Infantile spasms (HP:0012469)*, with all other phenotypes in the HPO using the FQ-HP and E-HP embedding methods. Points marked in black represent the cosine similarity values with phenotypes that are descendants of the reference phenotype. Points marked in grey are the cosine similarity values of non-descendant phenotypes.

## 5 Supplementary Method S3

We formally define Info Score as,

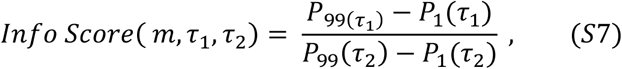

where *m* is the main phenotype, and *P*_*k*_(*τ*) is the *k*^*th*^ percentile of the pair-wise similarity values between *m* and all other phenotypes using the embedding technique *τ*. See Y and X in **Figure 4** representing the numerator and denominator of **Equation S7** with *τ*_1_ = FQ-HP and *τ*_2_ = E-HP, respectively. If the Info Score is very close to 1, there is no significant change in the cosine similarity values using the method *τ*_1_ over *τ*_2_. However, as an Info Score moves toward values smaller or greater than 1, it shows a significant change in the cosine similarity values and vector representations of the phenotypes.

## 6 Supplementary Table S1

**Table S1.**
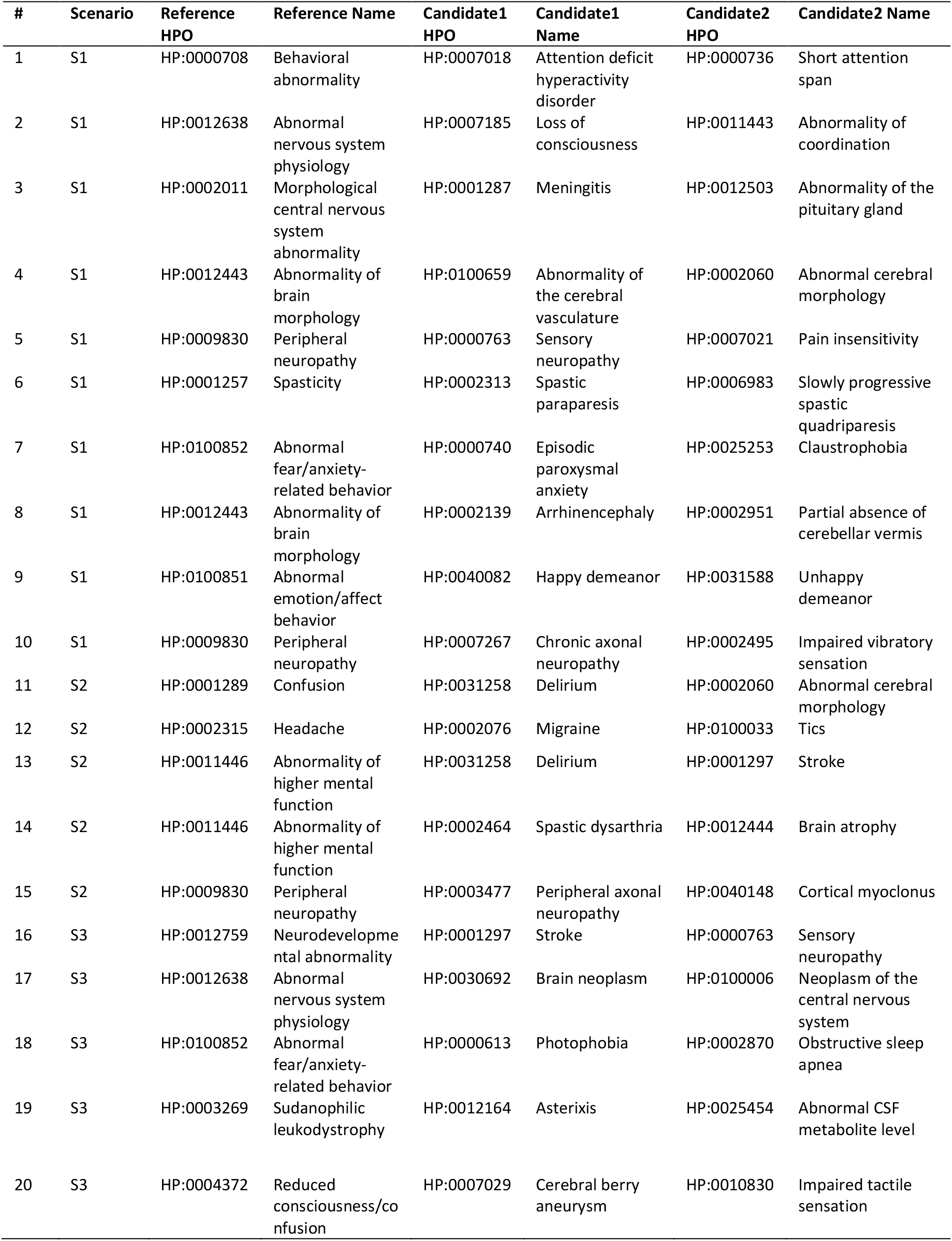
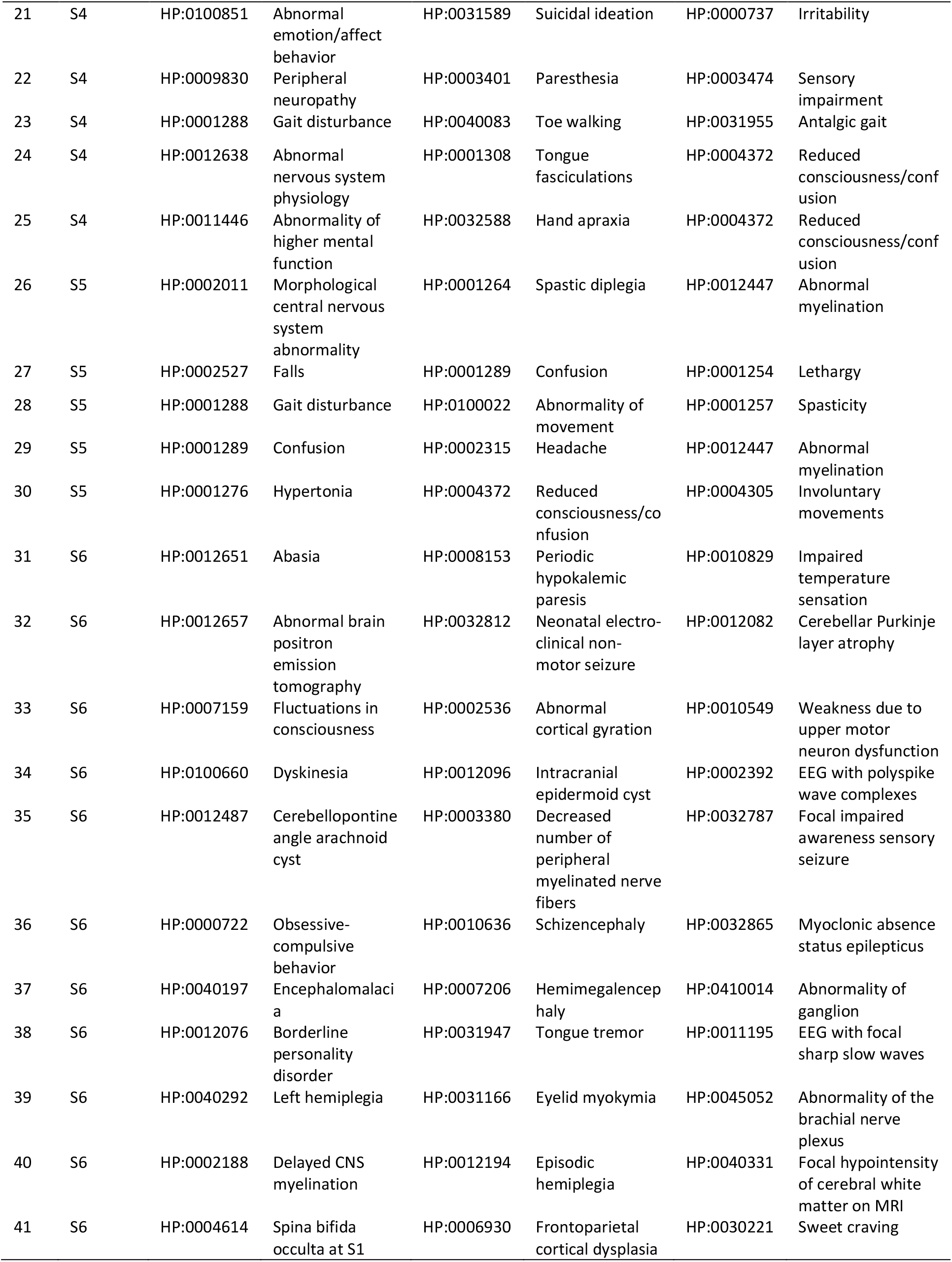
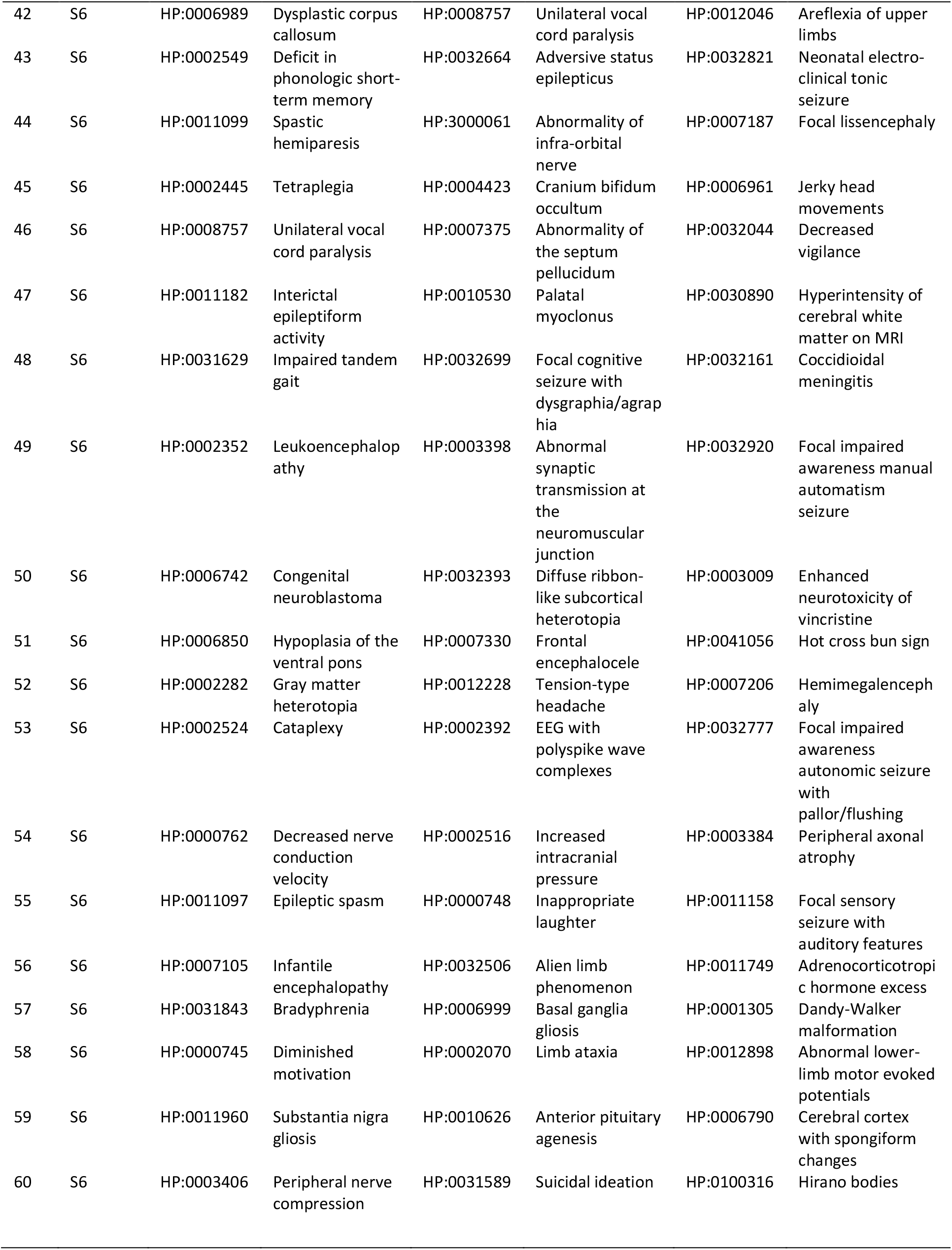
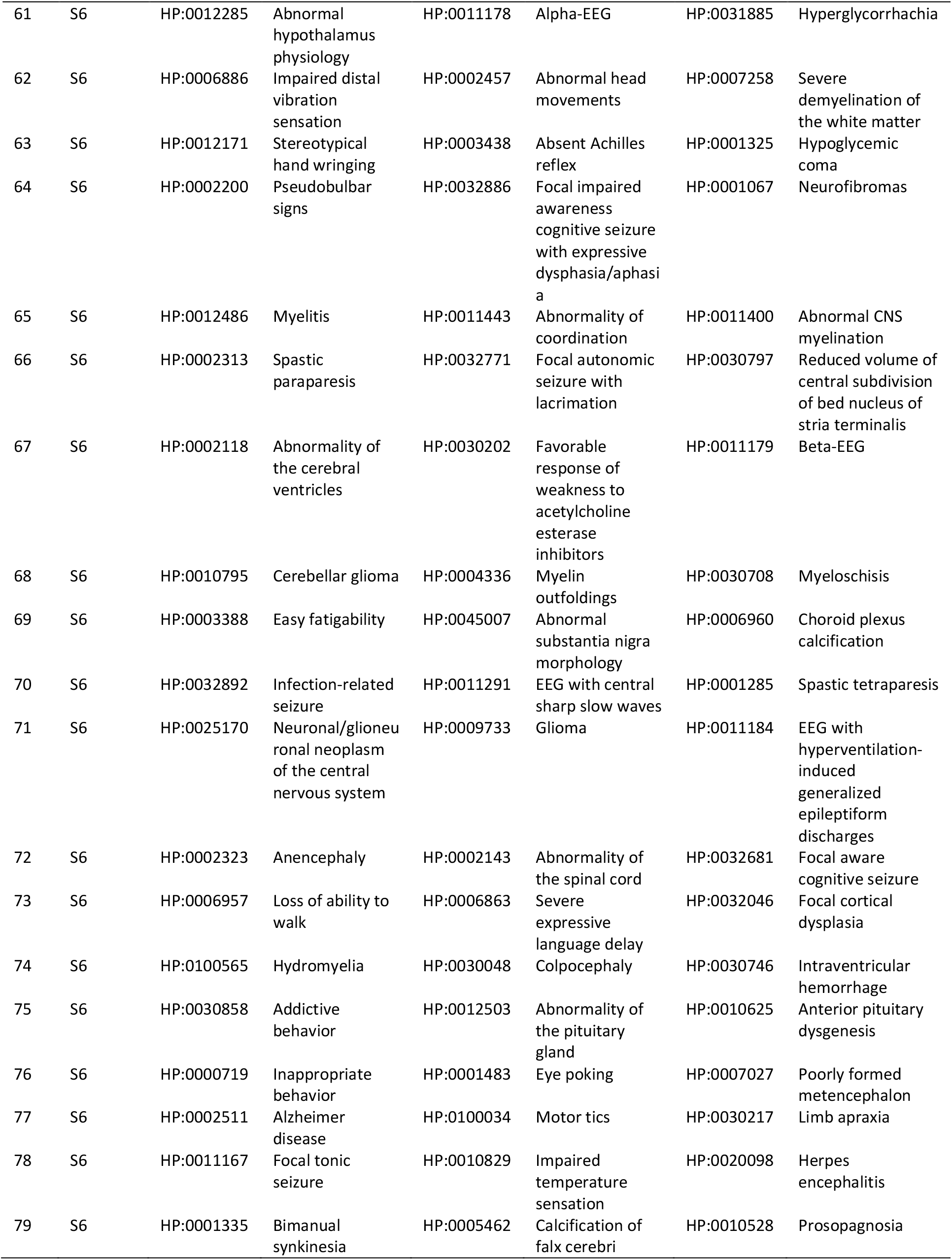
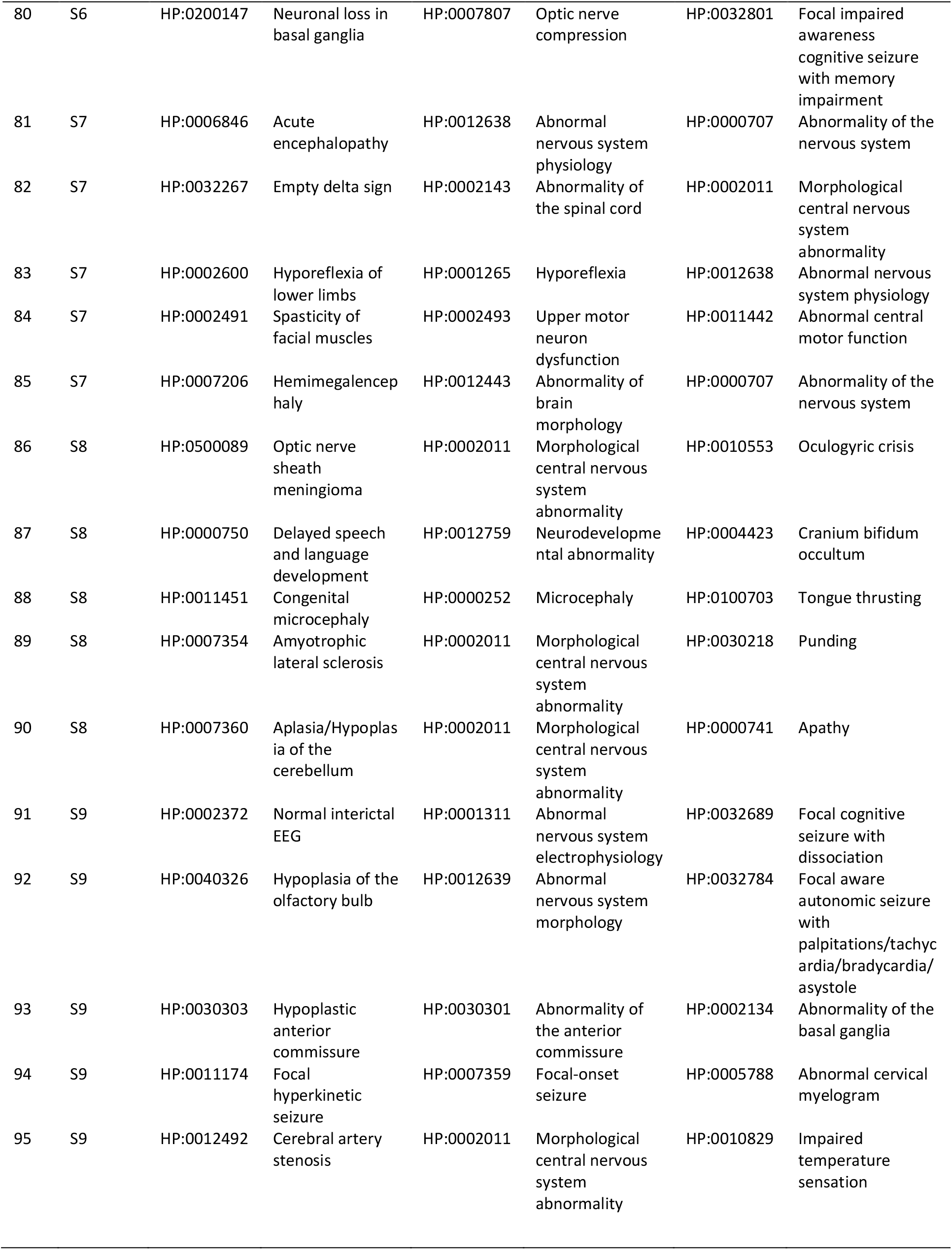
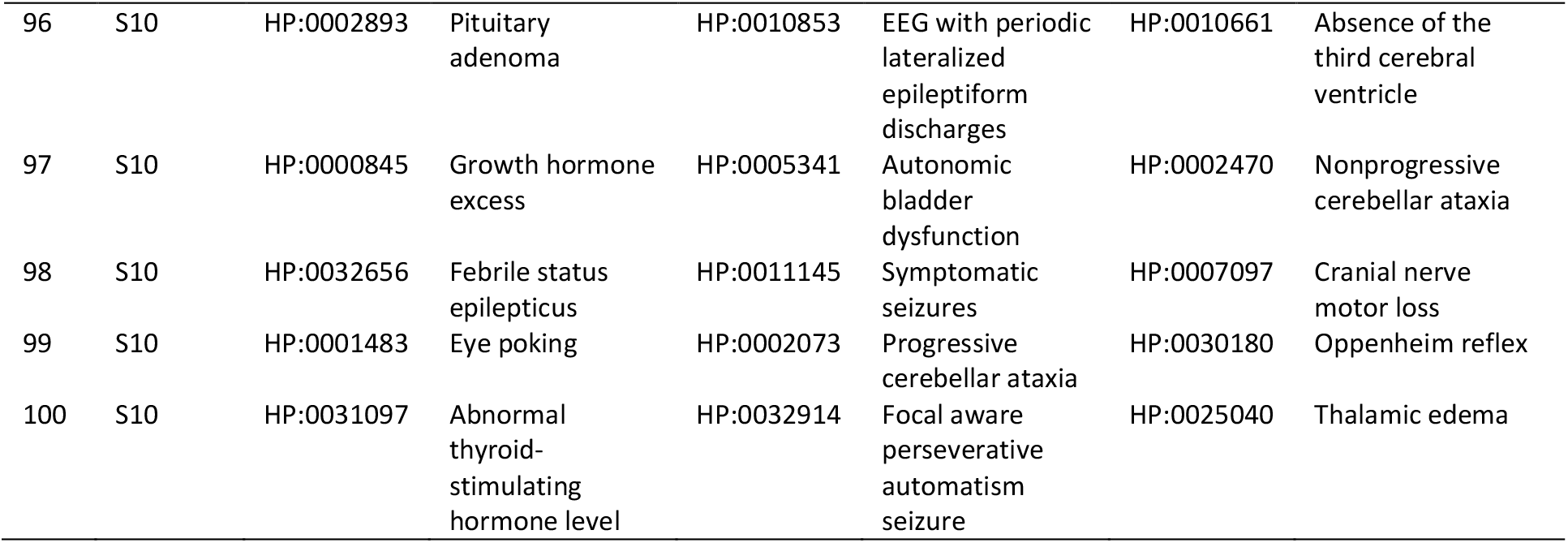
List of the 100 trios used for evaluation against expert opinion.

## 7 Supplementary Method S4

### 7.1 Agreement level calculation

Referring to how we generated the dataset, each prioritization could indicate candidate 1, candidate 2, or none (in case of uncertainty or tie). Thus, the minimum/chance level agreement is when only 5 out of the 13 experts (38.46%) have the same choice. When 6, 7, or 8 experts (≤61.53%) agree, we call it a “fair-level” agreement. In the case of an agreement between 9 or 10 experts (≤76.92%), we categorize the agreement level as “substantial”. Finally, it is called a “high-level” agreement if more than 11 experts (>76.92%) agree. **Figure 5B** (in the main manuscript) displays the distribution of agreement levels for the 100 trios.

## Footnotes

1 Also a tie between embedding techniques and Resnik

## References

1 Jha, A.K., et al., Use of electronic health records in US hospitals. New England Journal of Medicine, 2009. 360(16): p. 1628–1638.

2 Evans, R.S., Electronic health records: then, now, and in the future. Yearbook of medical informatics, 2016. 25(S 01): p. S48–S61.

3 Weng, C., N.H. Shah, and G. Hripcsak, Deep phenotyping: embracing complexity and temporality—towards scalability, portability, and interoperability. Journal of biomedical informatics, 2020. 105: p. 103433.

4 Kohler, S., et al., The Human Phenotype Ontology project: linking molecular biology and disease through phenotype data. Nucleic Acids Res, 2014. 42(Database issue): p. D966–74.

5 Kohler, S., et al., The Human Phenotype Ontology in 2017. Nucleic Acids Res, 2017. 45(D1): p. D865–D876.

6 Lewis-Smith, D., et al., Modeling seizures in the Human Phenotype Ontology according to contemporary ILAE concepts makes big phenotypic data tractable. Epilepsia, 2021. 62(6): p. 1293–1305.

7 Kohler, S., et al., The Human Phenotype Ontology in 2021. Nucleic Acids Res, 2021. 49(D1): p. D1207–D1217.

8 Groza, T., et al., The human phenotype ontology: semantic unification of common and rare disease. The American Journal of Human Genetics, 2015. 97(1): p. 111–124.

9 Galer, P.D., et al., Semantic similarity analysis reveals robust gene-disease relationships in developmental and epileptic encephalopathies. The American Journal of Human Genetics, 2020. 107(4): p. 683–697.

10 Helbig, I., et al., A recurrent missense variant in AP2M1 impairs clathrin-mediated endocytosis and causes developmental and epileptic encephalopathy. The American Journal of Human Genetics, 2019. 104(6): p. 1060–1072.

11 Lewis-Smith, D., et al., Phenotypic homogeneity in childhood epilepsies evolves in gene-specific patterns across 3251 patient-years of clinical data. European Journal of Human Genetics, 2021. 29(11): p. 1690–1700.

12 Lewis-Smith, D., et al., Computational analysis of neurodevelopmental phenotypes—harmonization empowers clinical discovery. Human Mutation, 2022.

13 Dewey, F.E., et al., Inactivating Variants in ANGPTL4 and Risk of Coronary Artery Disease. N Engl J Med, 2016. 374(12): p. 1123–33.

14 Gusarova, V., et al., Genetic inactivation of ANGPTL4 improves glucose homeostasis and is associated with reduced risk of diabetes. Nat Commun, 2018. 9(1): p. 2252.

15 Abul-Husn, N.S., et al., A Protein-Truncating HSD17B13 Variant and Protection from Chronic Liver Disease. N Engl J Med, 2018. 378(12): p. 1096–1106.

16 Köhler, S., et al., The human phenotype ontology in 2021. Nucleic acids research, 2021. 49(D1): p. D1207–D1217.

17 Savova, G.K., et al., Mayo clinical Text Analysis and Knowledge Extraction System (cTAKES): architecture, component evaluation and applications. Journal of the American Medical Informatics Association, 2010. 17(5): p. 507–513.

18 Deisseroth, C.A., et al., ClinPhen extracts and prioritizes patient phenotypes directly from medical records to expedite genetic disease diagnosis. Genetics in Medicine, 2019. 21(7): p. 1585–1593.

19 Aronson, A.R. and F.-M. Lang, An overview of MetaMap: historical perspective and recent advances. Journal of the American Medical Informatics Association, 2010. 17(3): p. 229–236.

20 Rodríguez-González, A., et al., Extracting diagnostic knowledge from MedLine Plus: a comparison between MetaMap and cTAKES Approaches. Current Bioinformatics, 2018. 13(6): p. 573–582.

21 Shivade, C., et al., A review of approaches to identifying patient phenotype cohorts using electronic health records. Journal of the American Medical Informatics Association, 2014. 21(2): p. 221–230.

22 Resnik, P., Using information content to evaluate semantic similarity in a taxonomy. arXiv preprint cmp-lg/9511007, 1995.

23 Li, B., et al., Effectively integrating information content and structural relationship to improve the GO-based similarity measure between proteins. arXiv preprint 1001.0958, 2010.

24 Pesquita, C., et al. Evaluating GO-based semantic similarity measures. In Proc. 10th Annual Bio-Ontologies Meeting. 2007. Citeseer.

25 Helbig, I., et al., A Recurrent Missense Variant in AP2M1 Impairs Clathrin-Mediated Endocytosis and Causes Developmental and Epileptic Encephalopathy. Am J Hum Genet, 2019. 104(6): p. 1060–1072.

26 Lewis-Smith, D., et al., Computational analysis of neurodevelopmental phenotypes: Harmonization empowers clinical discovery. Hum Mutat, 2022.

27 Bengio, Y., A. Courville, and P. Vincent, Representation learning: A review and new perspectives. IEEE transactions on pattern analysis and machine intelligence, 2013. 35(8): p. 1798–1828.

28 Le, Q. and T. Mikolov. Distributed representations of sentences and documents. In International conference on machine learning. 2014. PMLR.

29 Mikolov, T., et al., Efficient estimation of word representations in vector space. arXiv preprint 1301.3781, 2013.

30 Grover, A. and J. Leskovec. node2vec: Scalable feature learning for networks. In Proceedings of the 22nd ACM SIGKDD international conference on Knowledge discovery and data mining. 2016.

31 Narayanan, A., et al., graph2vec: Learning distributed representations of graphs. arXiv preprint 1707.05005, 2017.

32 Shen, F., et al. Constructing node embeddings for human phenotype ontology to assist phenotypic similarity measurement. In 2018 IEEE International Conference on Healthcare Informatics Workshop (ICHI-W). 2018. IEEE.

33 Shen, F., et al., HPO2Vec+: Leveraging heterogeneous knowledge resources to enrich node embeddings for the Human Phenotype Ontology. Journal of biomedical informatics, 2019. 96: p. 103246.

34 Arcus Data Repository Team, Deidentified Arcus Data Repository, Version 1.4.4. Extracted: 2021/07/09: Arcus at Children’s Hospital of Philadelphia.

35 Šuster, S., S. Tulkens, and W. Daelemans, A short review of ethical challenges in clinical natural language processing. arXiv preprint 1703.10090, 2017.

36 Straw, I. and C. Callison-Burch, Artificial Intelligence in mental health and the biases of language based models. PloS one, 2020. 15(12): p. e0240376.

37 Thayer, J., J.M. Miller, and J.W. Pennington. Fault-Tolerant, Distributed, and Scalable Natural Language Processing with cTAKES. In AMIA. 2019.

38 Masanz, J.J. and S. Finan. CTAKES 4.0. 2021 [cited 2022; Available from: https://cwiki.apache.org/confluence/display/CTAKES/cTAKES+4.0.

39 Bodenreider, O., The unified medical language system (UMLS): integrating biomedical terminology. Nucleic acids research, 2004. 32(suppl_1): p. D267–D270.

40 Chapman, W.W., et al., A simple algorithm for identifying negated findings and diseases in discharge summaries. Journal of biomedical informatics, 2001. 34(5): p. 301–310.

41 Xian, J., et al., Assessing the landscape of STXBP1-related disorders in 534 individuals. Brain (accepted), 2021.

42 Lewis-Smith, D., et al., Phenotypic homogeneity in childhood epilepsies evolves in gene-specific patterns across 3251 patient-years of clinical data. Eur J Hum Genet, 2021.

43 Ganesan, S., et al., A longitudinal footprint of genetic epilepsies using automated electronic medical record interpretation. Genet Med, 2020.

44 Galer, P., et al., Semantic similarity analysis reveals robust gene-disease relationships in developmental and epileptic encephalopathies. Am J Hum Genet, 2020.

45 Crawford, K., et al., Computational analysis of 10,860 phenotypic annotations in individuals with SCN2A-related disorders. Genet Med, 2021. 23(7): p. 1263–1272.

46 Feng, Y., et al., The state of the art in semantic relatedness: a framework for comparison. The Knowledge Engineering Review, 2017. 32.

47 Harispe, S., et al., Semantic similarity from natural language and ontology analysis. Synthesis Lectures on Human Language Technologies, 2015. 8(1): p. 1–254.

48 Slimani, T., Description and evaluation of semantic similarity measures approaches. arXiv preprint 1310.8059, 2013.

49 Stevenson, M. and M.A. Greenwood. A semantic approach to IE pattern induction. In Proceedings of the 43rd Annual Meeting of the Association for Computational Linguistics (ACL’05). 2005.

50 Lambert, J., Statistics in brief: how to assess bias in clinical studies? 2011, Springer.

51 Perozzi, B., R. Al-Rfou, and S. Skiena. Deepwalk: Online learning of social representations. In Proceedings of the 20th ACM SIGKDD international conference on Knowledge discovery and data mining. 2014.

52 Tang, J., et al. Line: Large-scale information network embedding. In Proceedings of the 24th international conference on world wide web. 2015.

53 Liu, R., M. Hirn, and A. Krishnan, Accurately Modeling Biased Random Walks on Weighted Graphs Using $\textit {Node2vec+} $. arXiv preprint 2109.08031, 2021.

54 Pearson, K., LIII. On lines and planes of closest fit to systems of points in space. The London, Edinburgh, and Dublin philosophical magazine and journal of science, 1901. 2(11): p. 559–572.

55 Van der Maaten, L. and G. Hinton, Visualizing data using t-SNE. Journal of machine learning research, 2008. 9(11).

56 Molinaro, A.M., R. Simon, and R.M. Pfeiffer, Prediction error estimation: a comparison of resampling methods. Bioinformatics, 2005. 21(15): p. 3301–3307.

57 Ronald, A., et al., Phenotypic and genetic overlap between autistic traits at the extremes of the general population. Journal of the American Academy of Child & Adolescent Psychiatry, 2006. 45(10): p. 1206–1214.

58 Cogliati, F., F. Forzano, and S. Russo, Overlapping Phenotypes and Genetic Heterogeneity of Rare Neurodevelopmental Disorders. Frontiers in Neurology, 2021. 12.

## References (Supplemental)

1 Mikolov, T., et al., Efficient estimation of word representations in vector space. arXiv preprint 1301.3781, 2013.

2 Mikolov, T., et al., Distributed representations of words and phrases and their compositionality. Advances in neural information processing systems, 2013. 26.

3 Perozzi, B., R. Al-Rfou, and S. Skiena. Deepwalk: Online learning of social representations. In Proceedings of the 20th ACM SIGKDD international conference on Knowledge discovery and data mining. 2014.

